# A large-effect locus on chromosome 4 underlies circadian feeding rhythmicity in pigs

**DOI:** 10.64898/2026.09.01.748529

**Authors:** Wim Gorssen, Naveen Kumar Kadri, Xena Marie Mapel, Alexander S. Leonard, Qiongyu He, Carmen Winters, Negar Khayatzadeh, Claudia Kasper, Hubert Pausch

## Abstract

Circadian feeding rhythmicity reflects the distribution of feed intake across the daily 24-hour cycle. Regular feeding-fasting cycles play an important role in regulating metabolism and maintaining health, but endogenous and environmental factors can disrupt the normal circadian pattern of feed intake. Here, we quantified hourly feed intake in 3,470 Swiss Landrace and 15,181 Swiss Large White pigs using 40.5 million records from automated feeding systems. We used the proportion of days exhibiting a significant feeding rhythm after wavelet analysis to investigate variation in circadian feeding organization. Circadian feeding rhythmicity was influenced by sex and age, with higher rhythmicity observed in females and older animals. Variance components analysis revealed a strong additive genetic contribution and a high heritability (h² = 0.53-0.57). Pigs with higher circadian feeding rhythmicity had reduced nocturnal intake, improved feed conversion ratio, and lower day-to-day variability in feed intake, suggesting improved feed efficiency and resilience. Genome-wide association testing of 15.7 and 23.1 million imputed sequence variants identified a QTL on chromosome 4 explaining 5.3% and 2.9% of the phenotypic variance of circadian feeding rhythmicity in Swiss Landrace and Swiss Large White pigs, respectively. The QTL overlaps *OPRK1* and *NPBWR1*, two genes implicated in the regulation of feeding behavior, reward and energy homeostasis. Collectively, our findings establish circadian feeding behavior as a heritable trait in pigs and provide a basis to further investigate the genetic regulation of feeding rhythms in mammals and explore its use in animal breeding programs to improve feed efficiency and resilience.

**Significance Statement:** The timing of feed intake influences metabolism and health, but the genetic basis of feeding rhythms is poorly understood. Pigs are diurnal omnivores with daytime-dominated feeding patterns resembling those of humans, making them a useful model for studying circadian traits. Using over 40 million feeding records, we show that individual differences in 24-h feeding rhythmicity are highly heritable. Pigs with higher rhythmicity eat less at night and convert feed into growth more efficiently. A genomic region on porcine chromosome 4 near two candidate genes, *OPRK1* and *NPBWR1*, explains a substantial proportion of variation in feeding rhythmicity. These findings suggest that feeding rhythmicity may be a useful target trait for improving feed efficiency and resilience in livestock via selective breeding.

## Introduction

The circadian organization of feeding is coordinated by multiple endogenous oscillatory systems. The suprachiasmatic nucleus (SCN) of the hypothalamus serves as the principal circadian pacemaker synchronized by the light–dark cycle and plays a major role in organizing daily feeding rhythms, while feeding-related rhythms can also be synchronized by meal timing through circadian clocks outside the SCN (1–3). Circadian feeding patterns likely emerge from interactions between the molecular circadian clock and neuromodulatory pathways regulating feeding motivation and energy balance (4,5). As diurnal omnivores, pigs have a bimodal feeding pattern with two pronounced feeding bouts during the day and minimal intake at night (6), which is similar to humans (7). Consequently, pigs provide a valuable translational model for investigating how circadian organization of feeding relates to metabolism and health (8).

The adoption of automated feeding stations in pig farming enables precise, longitudinal assessment of feeding behavior in large cohorts of animals (9,10). Such data have revealed substantial inter-individual variation in circadian feeding rhythmicity, with rhythmicity generally increasing with age (10). A recent study reported that circadian feeding rhythmicity is moderately heritable in French Large White pigs at the end of the finishing period (h^2^=0.35) (11). However, genomic loci underlying this variation have not been identified.

Perturbations of circadian rhythms are associated with metabolic disorders in humans, pigs and other mammals (1,10) and feeding during low-activity periods has negative metabolic consequences across species (12,13). For instance, night feeding in pigs increases fat deposition and reduces feed efficiency (14,15). In a divergent selection experiment, the more feed-efficient line showed higher feeding rhythmicity and allele frequency shifts around core circadian clock genes (16), suggesting a biological relationship between circadian feeding rhythmicity and metabolic efficiency.

Here, we leverage 40.5 million feeding records to characterize circadian feeding rhythmicity and dissect its genetic basis in 18,651 pigs. We quantify the heritability of feeding rhythmicity, assess its relationship with production traits, and identify genomic loci associated with variation in circadian feeding rhythmicity using imputed whole-genome sequence variant genotypes. Our findings reveal a major QTL for circadian feeding rhythmicity in pigs and connect variation in the temporal organization of feeding with feed efficiency and feeding stability.

## Results

### Circadian feeding rhythmicity in pigs is a heritable age- and sex-dependent trait

We analyzed 40,482,476 feeding events from 3,470 Swiss Landrace (SLR) and 15,181 Swiss Large White (SLW) pigs between 75 and 155 days of age to characterize hourly feed intake under ad libitum feeding conditions. Data were recorded by automated feeding stations in an experimental breeding farm from 2017 to 2026. A subset of the phenotyped cohort (1,403 SLR and 2,593 SLW pigs) had microarray-derived genotypes at 57,528 single nucleotide polymorphisms (SNPs). Using wavelet-based analyses of hourly feed intake, we quantified the proportion of days exhibiting a significant ∼24-h feeding pattern (PropCirc; Fig. 1). Hence, PropCirc is an operational measure of circadian feeding rhythmicity.

**Figure 1.**
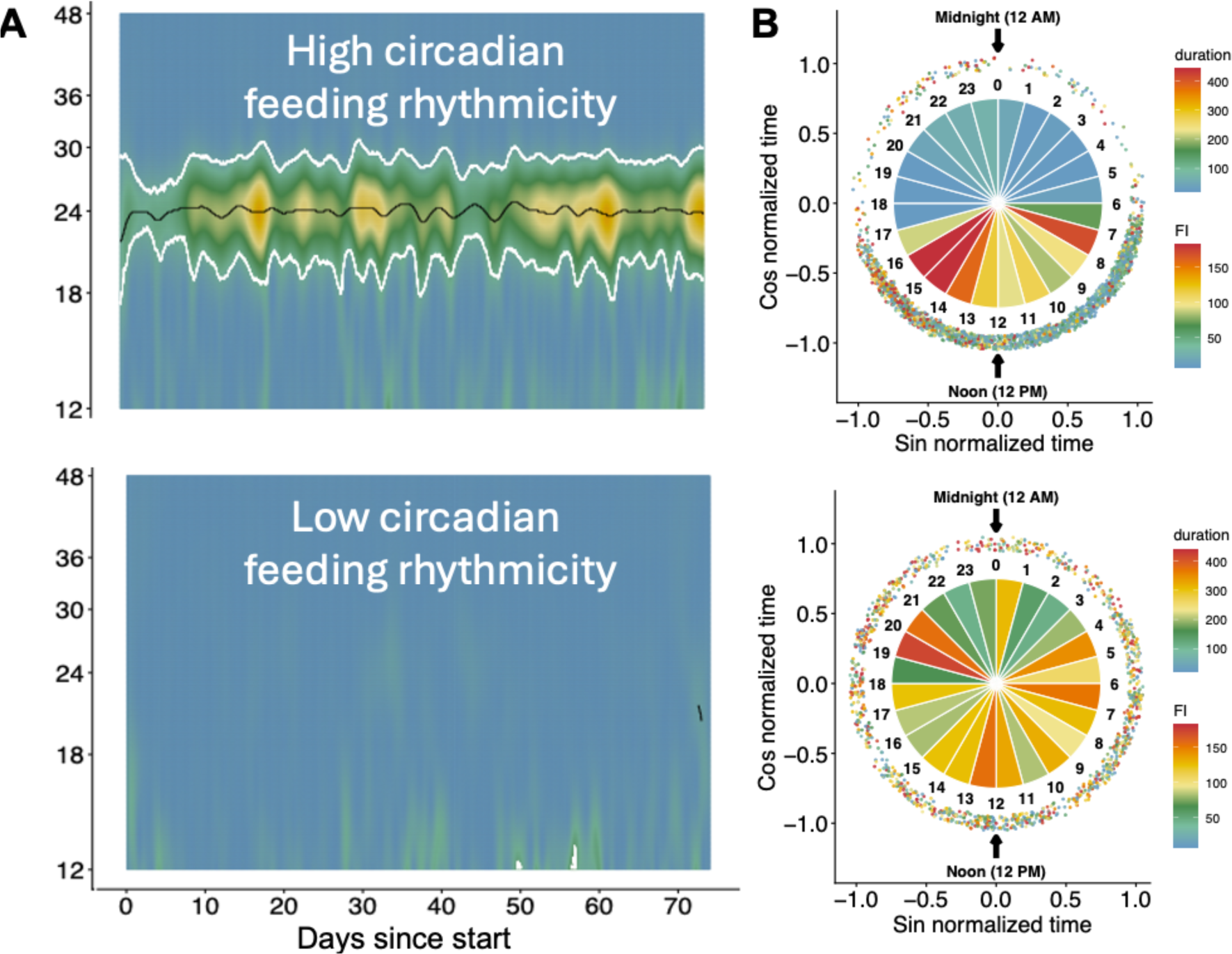
Feeding pattern over a period of 81 days in two exemplary pigs from 75 to 155 days of age with high (top) and low (bottom) circadian feeding rhythmicity. **A**: Wavelet spectra showing periodicity of hourly feed intake across the finishing period. Warmer colors indicate higher periodicity, and white contours indicate significant rhythmicity (P < 0.05). **B**: Circular 24-h plots summarizing feeding behavior. Pie segments indicate mean feed intake (FI) per hour (g), and dots represent the duration of feeding visits (s). Pigs with higher circadian feeding rhythmicity show consolidated daytime feeding and low nocturnal feeding, whereas pigs with lower rhythmicity exhibit feeding activity distributed more evenly across the 24-h cycle.

Circadian feeding rhythmicity showed substantial inter-individual variation in both breeds (Fig. 2). Mean PropCirc (%) was higher in SLR (55.6 ± 28.7%) than in SLW (43.8 ± 30.4%), but this difference was not statistically significant (P = 0.37). Females exhibited significantly higher circadian feeding rhythmicity than males (+9.4 percentage points, P = 7.5 × 10^-19^) and castrates (+18.9 percentage points, P = 1.3 × 10^-216^) across both breeds (Fig. 2). PropCirc increased significantly (+4.6 percentage points) from the early growing period (75-102 days of age) to the late finishing period (131-155 days of age; P = 6.0 × 10^−8^; Fig. S1).

**Figure 2.**
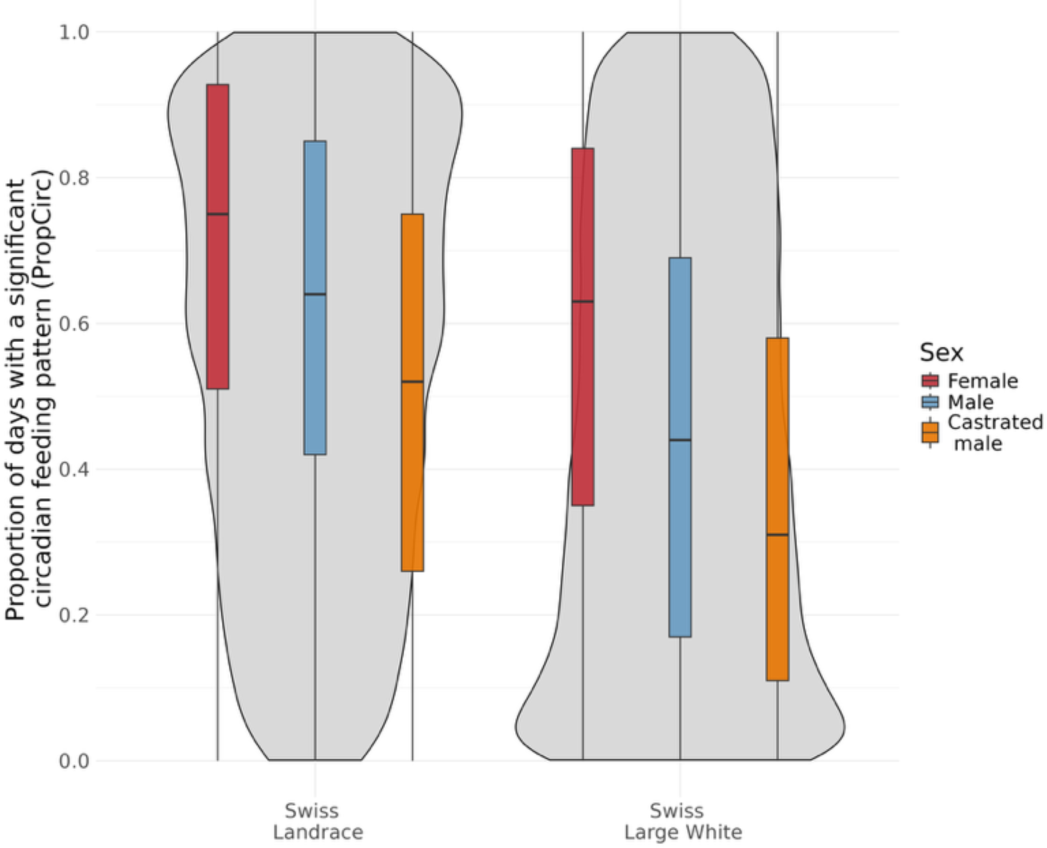
Distribution of circadian feeding rhythmicity across breeds and sex classes. Swiss Landrace shows higher mean PropCirc than Swiss Large White (55.6% vs 43.8%). Females exhibit consistently higher PropCirc than males and castrates across both breeds.

The narrow-sense heritability of PropCirc was high (h² = 0.53-0.57) in both breeds, indicating that additive genetic effects explain more than half of the observed phenotypic variation (Fig. S2; Tables S1-S2). In contrast, common environmental (pen) effects were modest (SLR: 9.2%; SLW: 7.7%) (Tables S1-S2) and non-additive genetic variance was negligible in both breeds.

### Circadian feeding rhythmicity is associated with nocturnal intake and feed efficiency

Higher circadian feeding rhythmicity was associated with a lower proportion of feed intake during the night in both breeds, with strong phenotypic (SLR: r = −0.53; SLW: r = −0.54) and genetic correlations (SLR: r_g_ = −0.63; SLW: r_g_ = −0.67; Fig. S2 and Tables S3–S4). Higher PropCirc was also phenotypically associated with a lower feed conversion ratio (FCR) in both breeds (SLR: r = - 0.14; SLW: r = −0.25), indicating improved feed efficiency. Genetic correlations with FCR were small and negative (SLR: r_g_ = −0.13; SLW: r_g_ = −0.09), although the 95% HPD interval included zero in SLR. PropCirc showed little relationship with average daily gain at either the phenotypic (SLR: r = 0.03; SLW: r = 0.01) or genetic level (SLR: r_g_ = 0.02; SLW: r_g_ = −0.06).

PropCirc was positively associated with lean meat content in both breeds (phenotypic: r = 0.21 to 0.33; genetic: r_g_ = 0.11 to 0.17). Higher PropCirc was also associated with lower day-to-day variability in feed intake, measured as lnvar_FI_ (phenotypic: r = −0.24 to −0.23; genetic: r_g_ = −0.33 to - 0.24). Detailed genetic and phenotypic correlations among traits are provided in Fig. S2 and Tables S1–S4.

### A major QTL on chromosome 4 underlies circadian feeding rhythmicity

The high heritability of PropCirc suggests that this trait is amenable to genome-wide association testing. Using large whole-genome sequenced reference panels, we imputed array-derived genotypes of 1,403 SLR and 2,593 SLW pigs, that had phenotypes for PropCirc, to the sequence level. Based on power calculations following Goddard and Hayes (17), the SLR and SLW cohorts provide sufficient statistical power to identify loci explaining respectively at least 2% and 1% of the genetic variation of PropCirc. Genome-wide association analyses using imputed sequence variants identified a single pronounced association signal on porcine chromosome 4 for PropCirc that exceeded the Bonferroni-corrected significance threshold in both breeds (Fig. 3). Variants significantly associated with PropCirc resided within a ∼0.9 Mb interval with lead variants located at Chr4:77,597,154 (P = 1.7 × 10^-8^) in SLR and Chr4:77,487,163 (P = 4.5 × 10^-14^) in SLW (Dataset S1). The 0.9 Mb interval contains six protein-coding genes including *OPRK1* encoding opioid receptor kappa 1 and *NPBWR1* encoding neuropeptides B and W receptor 1, two genes implicated in feeding regulation and energy homeostasis in other mammals (18–20).

**Figure 3.**
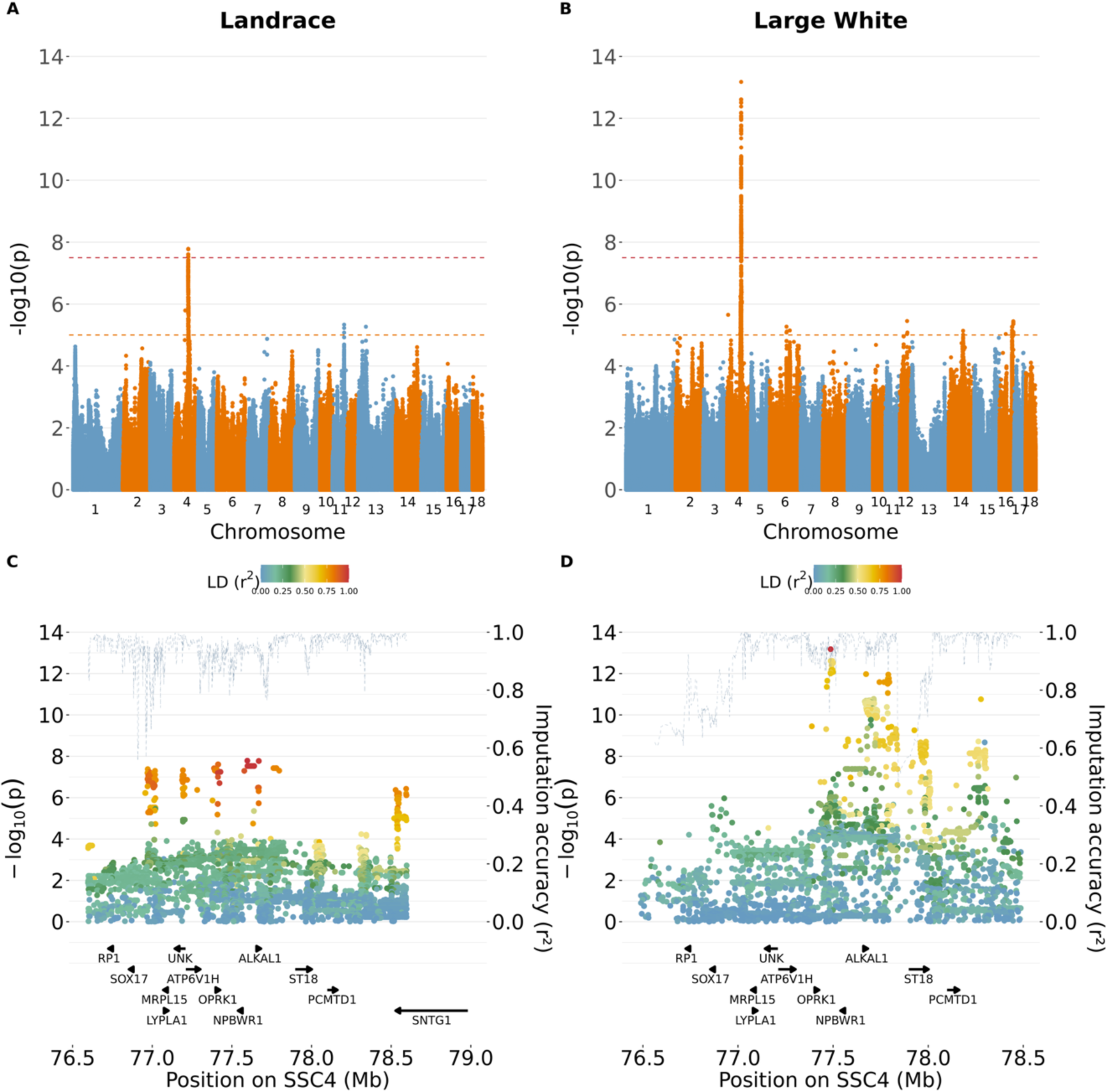
A quantitative trait locus on chromosome 4 is associated with circadian feeding rhythmicity. Manhattan plots representing the results of a genome-wide association study between imputed sequence variants and PropCirc in Swiss Landrace **(A)** and Swiss Large White **(B)** pigs. Dashed lines indicate genome-wide (red) and suggestive (orange) significance thresholds. Regional locus zoom plot displaying association p-values for variants at the chromosome 4 QTL against their chromosomal positions in Swiss Landrace **(C)** and Swiss Large White **(D)** pigs. Colors indicate linkage disequilibrium (r²) with the lead variant, and the dashed black line represents imputation accuracy.

The lead variants at the QTL exhibited additive effects, increasing PropCirc by +6.8 percentage points (±1.1 percentage points) in SLR and +7.7 percentage points (±0.8 percentage points) in SLW per effect allele (Fig. S3). Genome partitioning of phenotypic variation indicated that a 5-Mb window centered on the lead variant on chromosome 4 QTL explained a substantial proportion of the phenotypic variance (2.9-5.3%) and the genetic variance (6.4-10.3%) for PropCirc in both breeds (Tables S5-S6). This suggests that this region acts as a major-effect QTL for PropCirc. The QTL was absent when the association analysis was conditioned on the lead variant in both breeds, supporting the presence of a single underlying QTL. Notably, the chromosome 4 region was not associated with any production or feeding trait investigated (Fig. S4-S13), suggesting that its association is specific to circadian feeding rhythmicity rather than mediated by correlated production or feeding traits.

The variants with the strongest associations with PropCirc were in intergenic and intronic regions (Dataset S2), suggesting that regulatory mechanisms contribute to the observed association signal. Two missense variants in *OPRK1* (Chr4:77,414,587 A>G; P = 2.6 × 10^-8^ in SLR; P = 3.4 × 10^-5^ in SLW) and *ST18* (Chr4:77,978,095 A>G; P = 0.20 in SLR; P = 3.4 × 10^-9^ in SLW) exceeded the Bonferroni-corrected significance threshold, but not in both breeds. However, the missense variant in *ST18* was orders of magnitude less significant than the respective lead variant.

To further characterize the chromosome 4 QTL, we investigated structural variation within the associated interval (75.5-79.5 Mb). Long-read sequencing of 18 SLW pigs identified one SV in perfect linkage disequilibrium with the lead haplotype in SLW pigs (Fig. S14): a 307-bp deletion at Chr4:77,452,166 located in the intergenic region between *OPRK1* and *NPBWR1* (Fig. S15). However, none of the SVs in LD (r>0.8) with the top haplotype overlapped genes, further corroborating that coding variants are less likely to underlie the QTL.

Cis-eQTL analysis of 22 expressed genes within ±1 Mb of the SLW lead variant in hypothalamic tissue from 48 SLW pigs did not identify significant associations after multiple-testing correction (Table S7). *NPBWR1* was insufficiently expressed for analysis (SI Appendix; Fig. S16).

## Discussion

Circadian feeding rhythmicity is a highly heritable behavioral trait in pigs, varying with sex and age and associated with nocturnal feeding, feed efficiency, and feed intake variability. We identified a major-effect locus on chromosome 4 that is associated with circadian feeding rhythmicity in two breeds and contains the plausible candidate genes *OPRK1* and *NPBWR1*. Together, these findings highlight circadian feeding rhythmicity as a potentially valuable trait for pig production and provide a basis for investigating the genetic regulation of feeding rhythms in mammals.

A particularly pronounced source of variation in circadian feeding rhythmicity is sex. Females exhibit higher circadian feeding rhythmicity than intact males and castrated males in both breeds. To the best of our knowledge, this is the first demonstration of a sex difference in circadian feeding rhythmicity in pigs. Similar sex differences have been reported in rodents (21) and humans (22). Experimental studies in rodents further demonstrate that estradiol promotes circadian rhythmicity and that gonadal hormone manipulation, including castration, alters circadian behavior (21,23,24). Together, these findings suggest that modulation of circadian rhythms by sex hormones may represent a conserved feature across mammals. Circadian feeding rhythmicity also increases with age during the growing–finishing period, consistent with previous findings in pigs (10,11). This pattern may reflect robust maturation of circadian regulation (25), although learned or socially reinforced meal timing could also contribute to the observed feeding rhythms (26).

Circadian feeding rhythmicity shows high heritability in both breeds (h² = 0.53-0.57), while common environmental and non-additive genetic effects contribute little to phenotypic variation. The heritability estimates from our study are at the upper end of those previously reported in French Large White pigs at the final finishing phase (h^2^ = 0.35) (11), possibly reflecting different trait definitions and the relatively uniform environmental conditions under which the animals in the present study were housed. Higher circadian feeding rhythmicity is associated with reduced nocturnal feed intake and improved feed efficiency without adverse effects on growth. Together with previous studies showing detrimental metabolic effects of circadian misalignment (12–15), these findings support the hypothesis that alignment of feeding behavior with endogenous circadian rhythms improves metabolic efficiency. However, because PropCirc was derived from feeding behavior alone, our data cannot distinguish variation in the central circadian system from other circadian mechanisms responsive to meal timing or from downstream behavioral and metabolic processes that shape feeding patterns. In addition, higher circadian feeding rhythmicity was associated with lower day-to-day feed intake variability, a distinct phenotype previously proposed as an indicator of resilience in pigs (27,28). This association suggests a potential link with resilience, although this interpretation remains tentative given the limited direct health phenotypes available in this study.

The chromosome 4 region spanning the association signal (75.5-79.5 Mb) explains up to 5.3% of the phenotypic variance in circadian feeding rhythmicity, exceeding the variance typically attributed to individual loci affecting complex production traits in pigs (29). The same genomic region is associated with PropCirc in two distinct breeds and the association signal disappears after conditioning on the respective lead variant supporting a single underlying QTL. The lead and most strongly associated variants are predominantly noncoding, making regulatory variation a plausible mechanism, although associated coding variants are also present within the region.

*Opioid Receptor Kappa 1* (*OPRK1*) and *Neuropeptide B and W Receptor 1* (*NPBWR1*) are plausible positional and functional candidate genes. Both genes are expressed in the central nervous system and are implicated in neuromodulatory pathways linking feeding behavior, reward, and energy homeostasis (18–20,30). Evidence from experimental studies supports their potential involvement in circadian feeding regulation: κ-opioid signaling influences feeding motivation and meal organization in rodents (19,31), while *NPBWR1*-deficient mice display hyperphagia and reduced energy expenditure (32). Moreover, *NPBWR1* shows circadian expression in the suprachiasmatic nucleus (33), and neuropeptide W administration suppresses dark-phase feeding in rats (34). Together, these findings are consistent with the hypothesis that variation at this locus influences the temporal organization of feeding behavior rather than overall feed intake per se. *OPRK1* is most highly expressed in the pig hypothalamus, and PigGTEx reports cis-eQTLs in other tissues (35). However, our hypothalamic cis-eQTL analysis did not identify significant associations between genetic variants and the expression of genes within the QTL interval. Given the limited sample size and our analysis being restricted to a single tissue, we cannot preclude that regulatory effects remained undetected due to limited statistical power. Thus, further research is warranted to elucidate the exact mechanism through which the QTL impacts the circadian feeding organization.

Overall, our findings establish circadian feeding rhythmicity as a highly heritable behavioral phenotype in pigs and identify a major-effect locus on chromosome 4 near *OPRK1* and *NPBWR1* contributing to its variation. These results identify circadian feeding rhythmicity as a promising target for improving feed efficiency and resilience in livestock while also establishing pigs as a valuable model for investigating the genetic regulation of feeding rhythms and metabolic organization in mammals.

## Materials and Methods

### Animal ethics statement

Pigs included in this study were kept in compliance with Swiss legislation. Phenotypic and genotypic data were obtained within the routine breeding program of the Swiss breeding organization SUISAG® (Sempach, Switzerland). The experiment from which the transcriptomic data originated was approved by the competent veterinary authority of the Canton of Fribourg (approval no. 2021_28_FR/34009) and conducted in accordance with Swiss animal welfare legislation.

### Animals and data collection

Data were collected from purebred Swiss Landrace (SLR) and Swiss Large White (SLW) pigs at the testing station of SUISAG between January 2017 and January 2026. SLW animals originated from either the dam or sire line, which share a common ancestral population but have been bred separately since 2002 (36). Given their relatively recent shared origin, the two lines were analyzed jointly as SLW (Fig. S17). Feeding behavior was recorded using automated single-animal feeding stations (FIRE®, Osborne Industries, Inc., Osborne, KS; Schauer®, Schauer Agrotronic GmbH, Austria) equipped with radio-frequency identification antennas and integrated load cells. For each visit, individual feed intake, duration, and timestamp were recorded.

Animals were housed under standardized commercial conditions. Male breeding candidates were kept in a separate unit comprising 16 pens across four compartments. Full siblings of breeding candidates, including females and castrates, were mixed and tested in separate units containing eight pens across 12 compartments and four pens across four compartments. Each pen (median of 10 pigs) was equipped with one feeder, and pigs had ad libitum access to feed and water, hay, and natural daylight through windows. A two-phase feeding regimen was applied during the growing-finishing period. Pigs received a diet containing 13.5 MJ digestible energy/kg and 15.0% crude protein up to approximately 60 kg body weight, after which crude protein content was reduced to 13.2%.

At entry into the testing station, mean age and body weight were 72.9 ± 6.9 days and 26.4 ± 5.0 kg for SLR and 72.4 ± 7.3 days and 26.7 ± 4.7 kg for SLW. At the end of testing, mean age and body weight were 158.6 ± 12.2 days and 108.6 ± 4.5 kg for SLR and 157.7 ± 13.2 days and 109.5 ± 4.0 kg for SLW. All medical treatments were recorded throughout the trial. Animals not selected for breeding were slaughtered, and carcass and meat-quality traits were measured.

The raw dataset comprised 6,637,824 feeding records from 4,026 SLR pigs and 36,348,935 records from 18,158 SLW pigs.

### Feeding data quality control and hourly aggregation

Quality control and statistical analyses were performed in R v4.4.0. Feeding records were filtered following procedures for automated pig feeding data described previously (37–39). Only pigs entering the animal performance testing station after 1 January 2017 were retained. Feed intake per visit was restricted to −20 to 3,000 g. For visits with a recorded duration of 0 s, intake was additionally required to be between −10 and 10 g. Visit start and end times were parsed in the Europe/Zurich time zone, and visit duration was recalculated from these timestamps. Negative time differences occurring on the autumn daylight-saving time transition were corrected by adding 1 h. Records with missing, negative, or >3,600-s durations were removed. Feeding rate was calculated as feed intake divided by visit duration and subjected to additional filters. For visits with intake >0 and <50 g, feeding rates >500 g/min were excluded. For visits with intake ≥50 g and duration <20 s, rates >600 g/min were excluded; visits with intake ≥50 g were additionally excluded when feeding rate exceeded 50 g/s (3,000 g/min). Visits with zero feeding rate were retained only when duration was ≤500 s, and nonzero feeding rates between −2 and 2 g/min were excluded. Finally, visits overlapping the preceding or following visit within the same feeding group were removed.

After record-level quality control, 6,228,348 feeding records from 3,923 SLR pigs and 34,254,128 records from 17,494 SLW pigs remained, corresponding to removals of 6.2% and 5.8% of the raw data, respectively. Feeding data were aggregated to hourly totals per pig. Hours without visits were assigned zero intake, yielding 24 hourly records per pig per day. For visits spanning multiple hours, intake was proportionally allocated according to the fraction of visit time within each hour. The first and last recording day per pig were removed to avoid incomplete days.

Only records between 75 and 155 days of age were retained. Animals were required to have over 15 days with feed intake records within this 81-day window. The resulting cohort comprised 3,470 SLR pigs with 6,097,237 hourly records and 15,181 SLW pigs with 26,858,081 hourly records for circadian analyses. Pedigree information was available for all animals.

### Quantification of circadian feeding rhythmicity and related phenotypes

Circadian feeding rhythmicity was quantified as the proportion of days showing a significant approximately 24-h feeding rhythm (PropCirc), following Bus et al. (10), with modifications to derive a single animal-level phenotype over the 75–155-day age interval. Wavelet analysis was used because it detects periodicity locally over time, allowing the presence of an approximately 24-h feeding rhythm to vary across the testing period. For each pig, hourly feed intake was detrended by fitting a LOESS curve (span = 0.75) across time and retaining the residuals. To account for changes in signal amplitude during growth, the residuals were divided, within consecutive 7-day windows, by the difference between the maximum and minimum residual within each window.

Continuous wavelet analysis was then applied to the normalized time series using the Morlet wavelet implemented in the WaveletComp package (40), evaluating periods between 8 and 48 h. Significance of wavelet power was assessed against 200 simulated white-noise time series. For each complete day, the median P value across the 24 hourly time points and wavelet periods between 23.5 and 24.5 h was calculated. Days with a median P ≤ 0.05 were classified as exhibiting a significant ∼24-h feeding rhythm. Days containing hourly observations flagged as missing were excluded, and PropCirc was calculated as the proportion of the remaining days exhibiting a significant ∼24-h rhythm. PropCirc therefore summarizes the persistence of detectable ∼24-h feeding rhythmicity across the testing period rather than assuming rhythmicity remains constant throughout growth.

The proportion of feed intake occurring during the night (FI_night_) was estimated from hourly intake profiles following Bus et al. (10). For each pig, a zero-adjusted gamma generalized additive model implemented in the gamlss package (41) modeled both the probability of feeding and the amount consumed, including a smooth trend across days and cyclic hourly patterns within consecutive 14-day periods. Only complete days were included, and periods containing fewer than 7 days were excluded. Predicted hourly feed intake was obtained by combining the probability of feeding with the predicted intake when feeding. FI_night_ was calculated as the proportion of predicted intake occurring between 21:00 and 05:00 h.

Production, feeding behavior, carcass composition, and resilience-related phenotypes were calculated to estimate phenotypic and genetic correlations with PropCirc. Average daily gain (ADG, kg/d) was calculated as the difference between body weight at the end and start of the testing period, divided by the duration of the period in days. Average daily feed intake (FI, kg/d), average daily time spent at the feeder (DUR, s/d), and average daily visit frequency (FREQ, visits/d) were obtained by summing the respective measures over the testing period and dividing by its duration in days. Feed conversion ratio (FCR, kg/kg) was calculated as FI divided by ADG.

A resilience indicator was estimated as the natural logarithm of the variance of the differences between day-to-day observed and predicted feed intake from linear regression (lnvar_FI_), following Gorssen et al. (42). For medical treatments, the total number of individual therapeutic interventions received by each pig during the finishing phase was summed (TRT), excluding group treatments and routine prophylactic procedures such as vaccinations, castration, and deworming.

Lean meat content (LMC, %) was measured using AutoFom III™ (Frontmatec, Smoerum A/S, Denmark). Intramuscular fat content (IMF) was measured using an NIR-Flex N-500 (Büchi, Flawil, Switzerland). Until December 2024, connective and adipose tissue were removed from loin samples before homogenization and NIR analysis; from January 2025 onward, IMF was measured directly on sliced loin samples without homogenization.

Phenotypic differences in PropCirc between breeds (SLR and SLW), sexes (female, intact male, and castrated male), and age classes (75-102, 103-130, and 131-155 days of age) were assessed using linear mixed-effects models implemented in the lme4 package in R (43). Models included breed and sex as fixed effects, body weight at arrival and number of feed intake records as covariates, farm of origin as an additional fixed effect, and testing group as a random intercept.

### Estimation of heritability and genetic correlations

Genetic parameters were estimated separately for each breed using Bayesian animal models implemented in gibbsf90+ software within a single-step genomic evaluation framework (44). Models were run for 500,000 samples, with a burn-in of 100,000 and a sampling interval of 1,000. Additive genetic effects were estimated using an animal model of the form

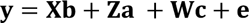

where **y** is the vector of phenotypes, **b** represents fixed effects and covariates, **a** represents additive genetic effects, **c** represents common pen effects, and **e** represents residual effects.

Models included sex (three levels: female, intact male, and castrated male) and farm of origin (11 levels for SLR and 28 levels for SLW) as fixed effects, and body weight at arrival and the number of days with available records during the evaluated period as covariates. Pen group was fitted as a random environmental effect (865 levels for SLR and 1,974 levels for SLW).

Additive genetic effects for each animal were modeled using a single-step H matrix (45–47), which integrates pedigree (**A**) and genomic (**G**) relationship information. Additive effects were assumed to follow a normal distribution with mean zero and variance 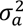 scaled by the H matrix: 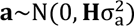. Common pen effects were assumed to follow a normal distribution with mean zero and variance 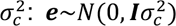; Residuals were assumed to follow a normal distribution with mean zero and variance 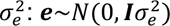; **X** and **Z** are incidence matrices for respectively fixed effects and random animal effects. The **H** matrix was constructed using the array-derived medium density genotypes.

Bivariate animal models were used to estimate genetic correlations between PropCirc and other traits. For each pair of traits, additive genetic effects were modeled jointly using the H matrix, and random pen effects were included for both traits. Posterior means and 95% highest posterior density intervals were reported.

### Genotypes, quality control, phasing and imputation

Microarray genotypes were provided by SUISAG, and quality control followed Gorssen et al. (48). The initial dataset contained 57,528 SNPs and 11,295 pigs (3,330 SLR; 7,965 SLW). SNP coordinates were aligned to the Sscrofa11.1 reference genome, alleles were updated, and duplicated positions were removed.

Using PLINK v1.9 (49), individuals with call rate <90% (49 SLR and 189 SLW), excess heterozygosity (>3 SD from the mean; 0 SLR and 0 SLW), or that did not pass sex checks (1 SLR and 3 SLW) were excluded. Duplicated animals identified using identity-by-descent estimates (--genome; PI_HAT >0.95) were removed (18 SLR and 154 SLW), as were animals showing Mendelian inconsistencies in the parentage check (--mendel; 11 SLW). SNPs with call rate <95% or minor allele frequency (MAF) <1% were removed. Autosomal SNPs deviating from Hardy– Weinberg equilibrium (P <0.0001) were excluded. For the X chromosome, heterozygous SNPs in males outside the pseudoautosomal region (0-7 Mb) were set to missing and subsequently treated as homozygous diploid. SNP-level QC was performed separately by breed.

After QC, the dataset contained 3,262 SLR pigs with 44,314 SNPs and 7,608 SLW pigs with 42,281 SNPs. Phenotypes were available for 1,403 genotyped SLR pigs and 2,593 genotyped SLW pigs. Population structure was assessed by principal component analysis in PLINK v1.9 (--pca; Fig. S17).

Array-derived genotypes were phased and sporadically missing genotypes imputed by breed using Beagle v5.4 (beagle.27Feb25.75f.jar) (50). Phased SLW genotypes were imputed to sequence level using an in-house reference panel of 122 sequenced SLW (≥10× coverage; mean = 15.9, SD = 5.0, maximum = 36.8) comprising 31,652,340 autosomal variants. Reference-based imputation with Beagle used an effective population size of 100, window size 10, overlap 2, and burn-in 5 (51). Sequence variants for SLR pigs were imputed using the SWine IMputation haplotype reference panel (52). Sex chromosomes were not imputed. After imputation, only variants with MAF > 0.01 and r^2^ > 0.5 were retained, yielding 15,689,634 variants (mean r^2^ = 0.902) for SLR and 23,134,253 variants (mean r^2^ = 0.914) for SLW.

### Genome-wide association study

Genome-wide association study (GWAS) analyses were performed in the 1,403 SLR and 2,593 SLW pigs with both PropCirc phenotypes and imputed genotypes for each breed separately using a linear mixed model-based framework implemented in GCTA v1.94.1 (53). Models included four principal components and body weight at arrival as quantitative covariates (--qcovar), and sex and farm of origin as categorical covariates (--covar), together with the genomic relationship matrix. To account for differences in the precision of aggregate phenotypes, residual variances were weighted according to the number of records contributing to each trait, using diagonal residual weights proportional to the inverse number of records via--reml-res-diag; weights were normalized to a mean of 1. Conditional association analyses were performed by fitting the respective lead variant as an additional fixed-effect covariate. Genome-wide significance was defined as P < 5×10^-8^. Manhattan plots were generated in R using the *manhattan* function from the qqman package (54).

Additive allelic effects and the phenotypic variance explained by individual lead variants were estimated using linear regression while controlling for the same covariates as in the GWAS model. Regional variance explained by the chromosome 4 QTL was estimated using multi-component genomic restricted maximum likelihood models in GCTA. For each breed, a 5-Mb window centered on the respective lead variant was defined as the candidate QTL region. The QTL window spanned Chr4:75,097,154–80,097,154 in SLR and Chr4:74,987,163–79,987,163 in SLW. Nineteen GRMs were fitted jointly, with one GRM for each autosome except chromosome 4, which was partitioned into the 5-Mb QTL region and the remainder of chromosome 4. Phenotypic variance explained was calculated as 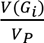, and genetic variance explained as 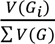 (Tables S5–S6).

To investigate whether dominance effects contributed to variation in PropCirc, we additionally performed non-additive association testing following He et al. (55). Non-additive effects refer here specifically to dominance deviations. Epistatic effects were not evaluated. Variants with MAF <0.05 were excluded to ensure sufficient representation of the homozygous genotype classes. Genotypes were recoded as RR = 0, RA = 2p, and AA = 4p − 2, where R and A denote the reference and alternative alleles and p is the alternative-allele frequency estimated within the respective GWAS cohort. The recoded genotypes were tested using GCTA with the same genomic relationship matrix and covariates as in the additive association analysis.

### Identification of candidate genes, functional annotation and structural variants

Genes located within ±1 Mb of genome-wide significant lead variants were identified using Ensembl annotations of the Sscrofa11.1 genome assembly. Functional consequences of significantly associated variants were predicted using Ensembl Variant Effect Predictor v114 (56). Positional candidate genes were evaluated based on gene annotation, predicted variant consequences, and published evidence for roles in feeding behavior, energy homeostasis, and circadian regulation.

Structural variation was evaluated within the identified 75.5-79.5 Mb region on chromosome 4 using short-read sequence data from 122 SLW pigs and PacBio HiFi sequence data from 18 SLW pigs. For the 18 HiFi-sequenced animals, high-molecular-weight DNA was extracted from frozen ear, tail, or blood tissue using the NEB Monarch® HMW DNA Extraction Kit. Sequencing was performed on the PacBio Sequel IIe platform using one SMRT cell per sample. HiFi reads were aligned to Sscrofa11.1 using pbmm2 v1.17 (57), followed by structural-variant calling with sawfish v1.0.1 (58).

Haplotype-based association analysis was performed on imputed sequence variants using an in-house Python script with sliding windows of 10 SNPs and a 5-SNP overlap. For the 122 short-read sequences, haplotypes at the most significantly associated locus were classified as homozygous carriers, heterozygous carriers, or non-carriers. Using Mosdepth (59), normalized coverage was calculated in 250-bp windows, and windows with absolute differences >3 SD were inspected using Integrative Genomics Viewer (IGV v2.19.5) (60).

### Hypothalamic gene expression and cis-eQTL analysis

Cis-eQTL analysis was performed using hypothalamic gene-expression and low-pass whole-genome sequencing data from 48 SLW pigs to investigate whether the chromosome 4 QTL was associated with gene expression. Hypothalamic tissue was prepared as described in Monney et al. (61). Samples were sequenced with the bulk RNA barcoding and sequencing approach (BRB-seq) (62) to estimate gene expression. DNA from the 48 samples was genotyped using low-pass whole-genome sequencing with subsequent genotype imputation by Gencove (Neogen, USA).

We considered all 29 annotated genes (13 protein-coding genes and 16 lncRNAs) located within ±1 Mb of the SLW lead GWAS variant at Chr4:77,487,163. The BRB-seq count matrix was filtered to retain genes with >6 counts in at least 10% of samples. After expression filtering, 22 genes were retained for cis-eQTL analysis. *NPBWR1* did not pass this filter because nonzero counts were detected in only five samples (maximum count = 5). *OPRK1* was low-to-moderately expressed (median counts = 15.5; SD = 12.1). We used edgeR (v4.8.2) (63) to normalize the filtered count matrix with trimmed mean of M-values (TMM).

For each retained gene, associations between gene expression and sequence variants in the chromosome 4 QTL region were tested using QTLtools v1.3.1 (64) (Table S7). DNA sequence variants with MAF <1% were excluded. We considered litter, age, and sex as covariates for the association testing, and included 17,919 variants within ±1 Mb of the transcription start site for *OPRK1*. We used the QTLtools *permute* function to perform 1,000 permutations and applied a 5% false discovery rate (FDR) to determine the significance threshold for the evaluated genes.

## Data availability

Individual-level feeding records, pedigree information, and genotype data are owned by SUISAG and cannot be made publicly available because of contractual and commercial restrictions. De-identified derived data underlying the principal findings of this study, including PropCirc phenotypes and nonidentifying metadata, GWAS summary statistics, data underlying the chromosome 4 QTL analyses, and the QTL expression and genotype data used for the cis-eQTL analysis, will be deposited in a public repository and made available upon publication. Analysis scripts required to reproduce the reported analyses will likewise be made publicly available upon publication. During peer review, these derived data and analysis scripts are available to editors and reviewers upon request. Access to the restricted primary data is controlled by SUISAG and is subject to approval by the data owner and applicable data-use agreements.

## Acknowledgments

We would like to thank all SUISAG employees and farmers that collected the data for this study. We acknowledge SUISAG for all their help and sharing their data. Thanks to Dr. Jacinta Bus for sharing the R-scripts for the wavelet-analysis to quantify circadian patterns of feed intake. We also thank Guy Maïkoff and the Agroscope piggery and abattoir teams for their support with animal handling and sample collection, and Paolo Silacci for laboratory work on the hypothalamic samples.

