## Supporting information for "A large-effect locus on chromosome 4 underlies circadian feeding rhythmicity in pigs"

### Supporting Information for
A large-effect locus on chromosome 4 underlies circadian feeding rhythmicity in pigs

Wim Gorssen[1]*, Naveen Kumar Kadri[1], Xena Marie Mapel[1], Alexander S. Leonard[1], Qiongyu He[1], Carmen Winters[2], Negar Khayatzadeh[3], Claudia Kasper[4] and Hubert Pausch[1]

[1]Animal Genomics, Department of Environmental Systems Science, Universitätstrasse 2, 8092 Zürich, Switzerland
[2]Independent researcher, Zürich, Switzerland
[3]SUISAG, Allmend 10, Sempach, 6204, Switzerland
[4]Agroscope, Animal GenoPhenomics, Tioleyre 4, 1725, Posieux, Switzerland

Wim Gorssen


**This PDF file includes:**

Supporting text
Figures S1 to S17
Tables S1 to S7
Legends for Datasets S1 to S2

**Other supporting materials for this manuscript include the following:**

Datasets S1 to S2

**Supporting Information Text**

**Supplementary results**

No significant cis-eQTLs were detected for the expressed protein-coding and long non-coding genes evaluated within ±1 Mb of the SLW PropCirc lead variant at Chr4:77,487,163 after correction for multiple testing (5% FDR; Table S7). No significant association with the SLW PropCirc lead variant itself was detected for any of the evaluated genes (Table S7).

The SLW PropCirc lead variant (Chr4:77,487,163; C/CACACACACACA) lies within a multiallelic AC tandem-repeat locus. PacBio HiFi long-read data independently confirmed repeat-length variation at this locus, including an allele corresponding to the GWAS indel as well as additional repeat-length alleles. However, this variation was poorly resolved in the low-pass sequencing data used for the eQTL cohort, limiting reliable evaluation of the lead GWAS variant as an expression-associated variant. In the low-pass sequencing data, variation at this locus was predominantly represented as C or CA alleles, whereas the CACACACACACA allele was observed in only one heterozygous individual and had uncertain imputation quality. Accordingly, the nominal association P values reported for the SLW lead variant in Table S7 should be interpreted cautiously.

The absence of significant cis-eQTLs may reflect limited statistical power due to the small sample size of 48 pigs, as well as limited resolution of the multiallelic tandem-repeat locus in the low-pass sequencing data. Thus, the present analysis does not provide evidence that the chromosome 4 PropCirc QTL acts through hypothalamic expression of the genes evaluated in this interval, but it also does not exclude a regulatory mechanism. Larger cohorts with higher-confidence sequence genotypes and gene-expression data will be required to resolve the regulatory consequences of the QTL.

**Figures**

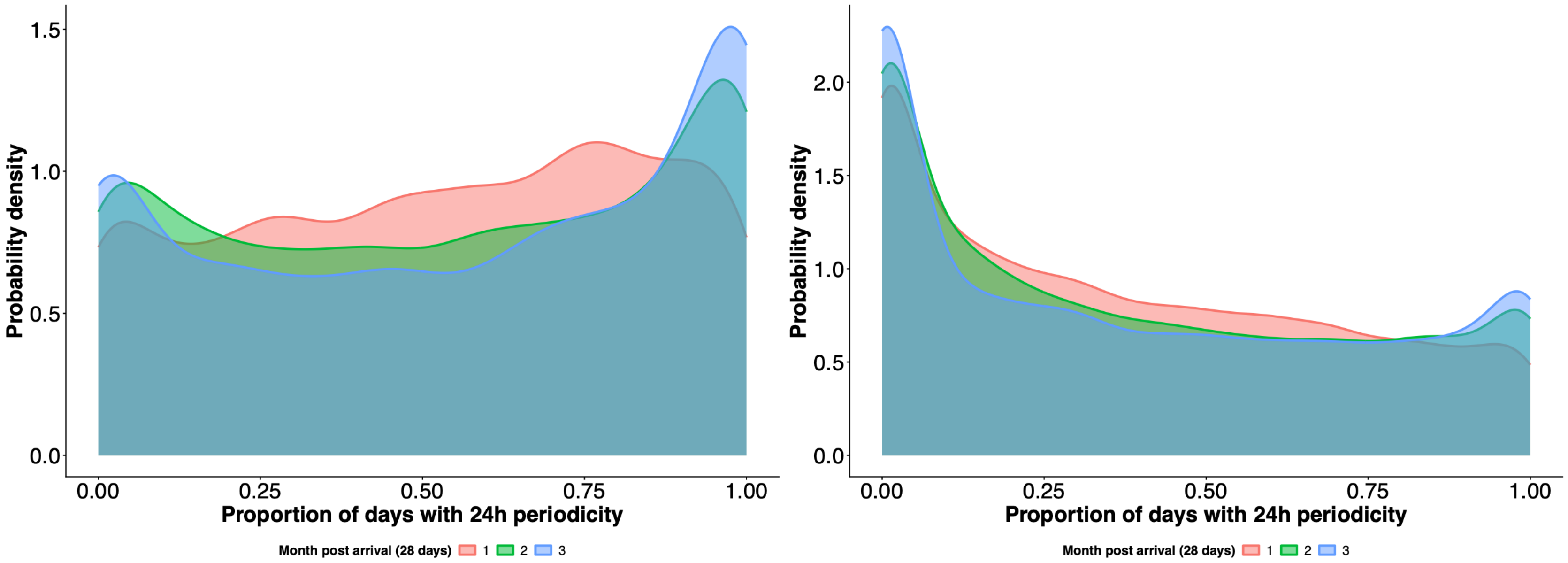


**Fig. S1. Age-related increase in circadian feeding rhythmicity.**

Distribution of the proportion of days with significant circadian feeding rhythm (PropCirc) across three age classes (75–102, 103–130, and 131–155 days). PropCirc increased with age, rising from 49 to 61% in SLR and from 41 to 46% in SLW between early and late finishing phases, with an overall effect of +4.6 percentage points ($P = 6.0 \times 10^{-8}$).

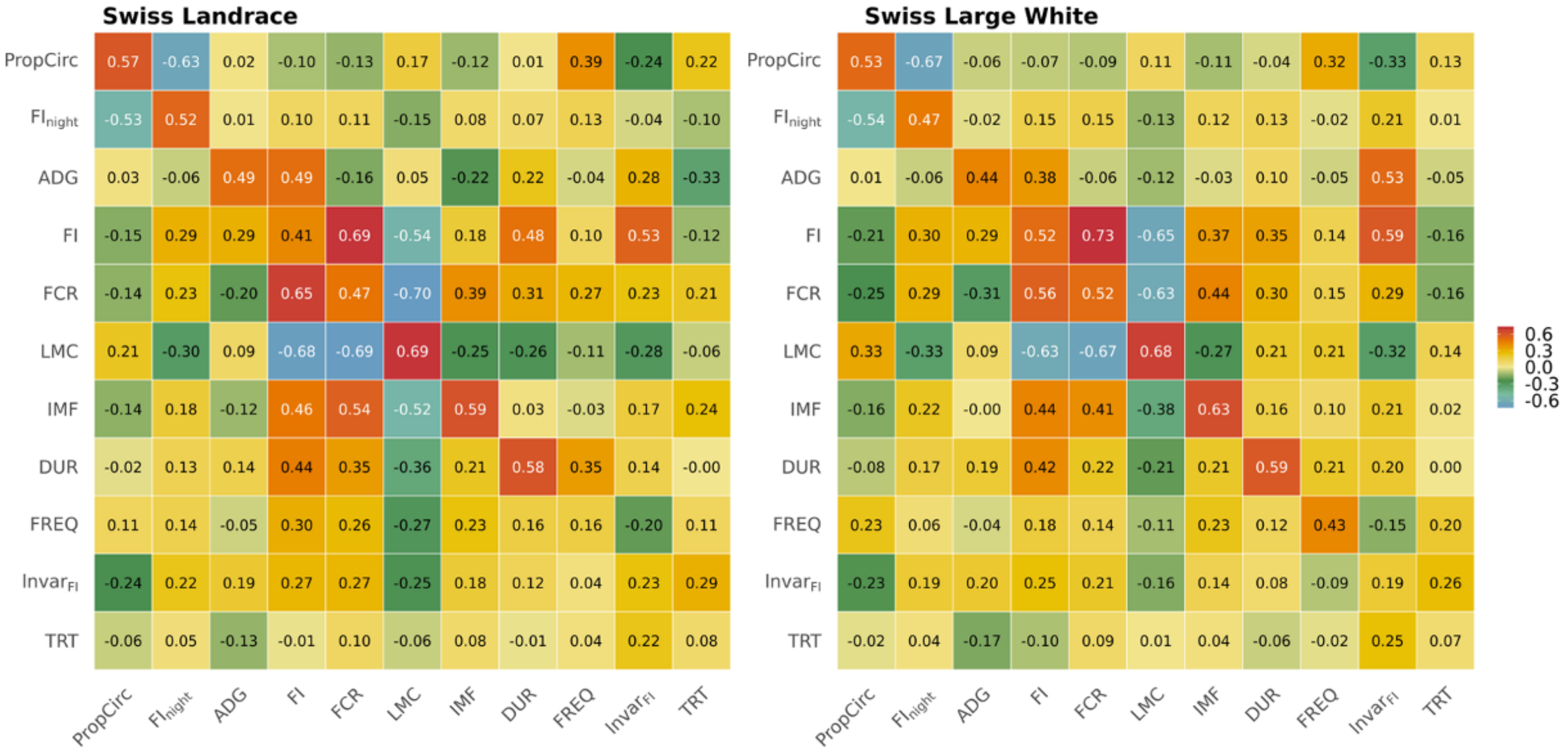


**Fig. S2. Genetic and phenotypic correlation structure of feeding and production traits.**

Heatmaps show genetic correlations (above diagonal), heritabilities (diagonal) and phenotypic correlations (below diagonal) per breed. Stronger circadian feeding rhythmicity (PropCirc) was associated with reduced nocturnal feeding, improved feed efficiency, and lower variability in feed intake. Detailed estimates and 95% highest posterior density intervals are provided in Tables S1-S4. Abbreviations: $FI_{night}$: Proportion of feed intake during night, ADG: Average daily gain, FI: Average daily feed intake, FCR: Feed conversion ratio, LMC: Lean meat content, DUR: Mean daily visit duration, FREQ: Mean daily visit frequency, $Invar_{FI}$: log variance of day-to-day feed-intake deviations, a previously proposed resilience indicator, TRT: number of medical treatments, IMF: intramuscular fat

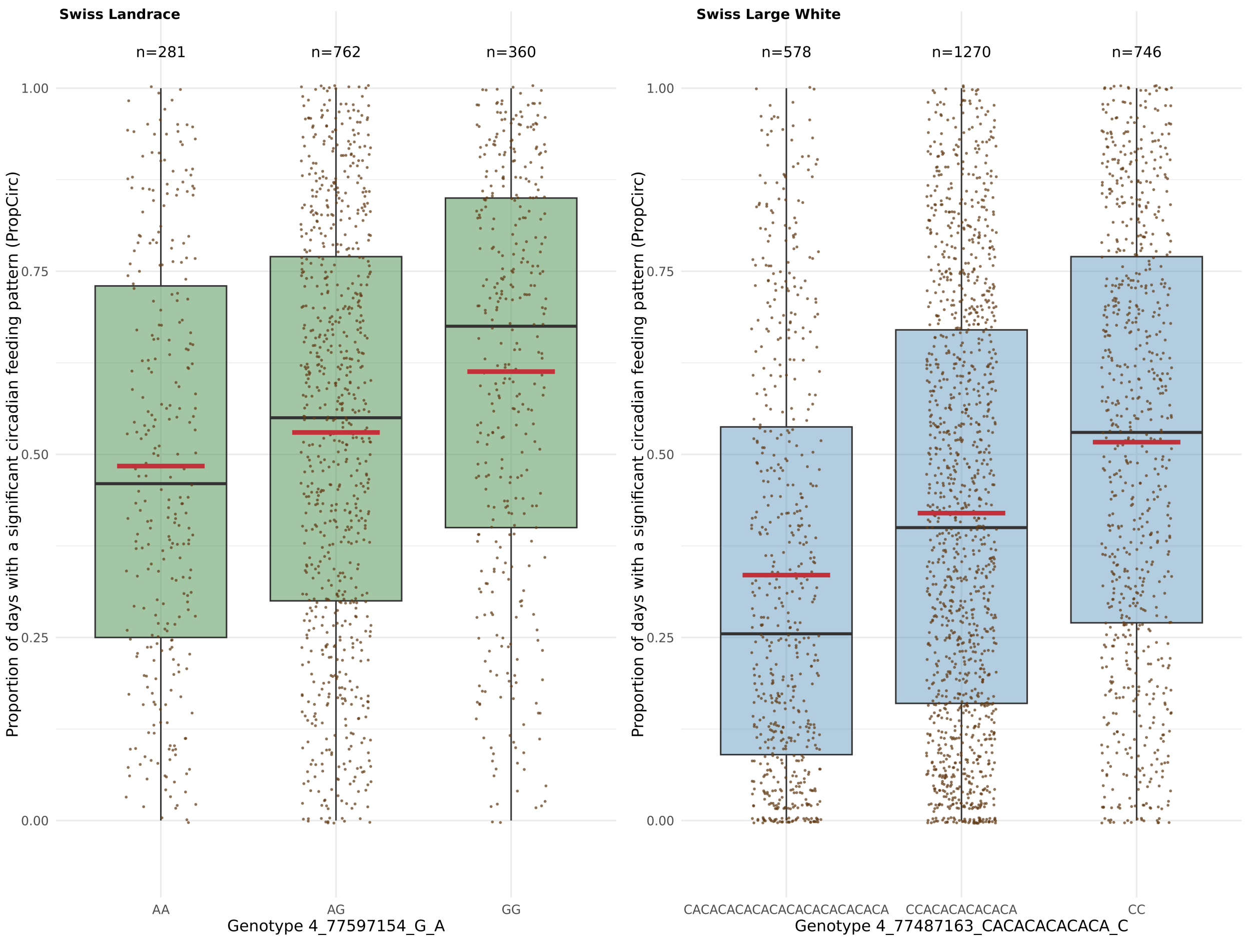


**Fig. S3. Effect of the lead SNP on circadian feeding rhythmicity.**

Boxplots showing the distribution of PropCirc by genotype at the lead SNP in Swiss Landrace (A) and Swiss Large White (B). Red lines indicate mean phenotypic values for each genotype class.

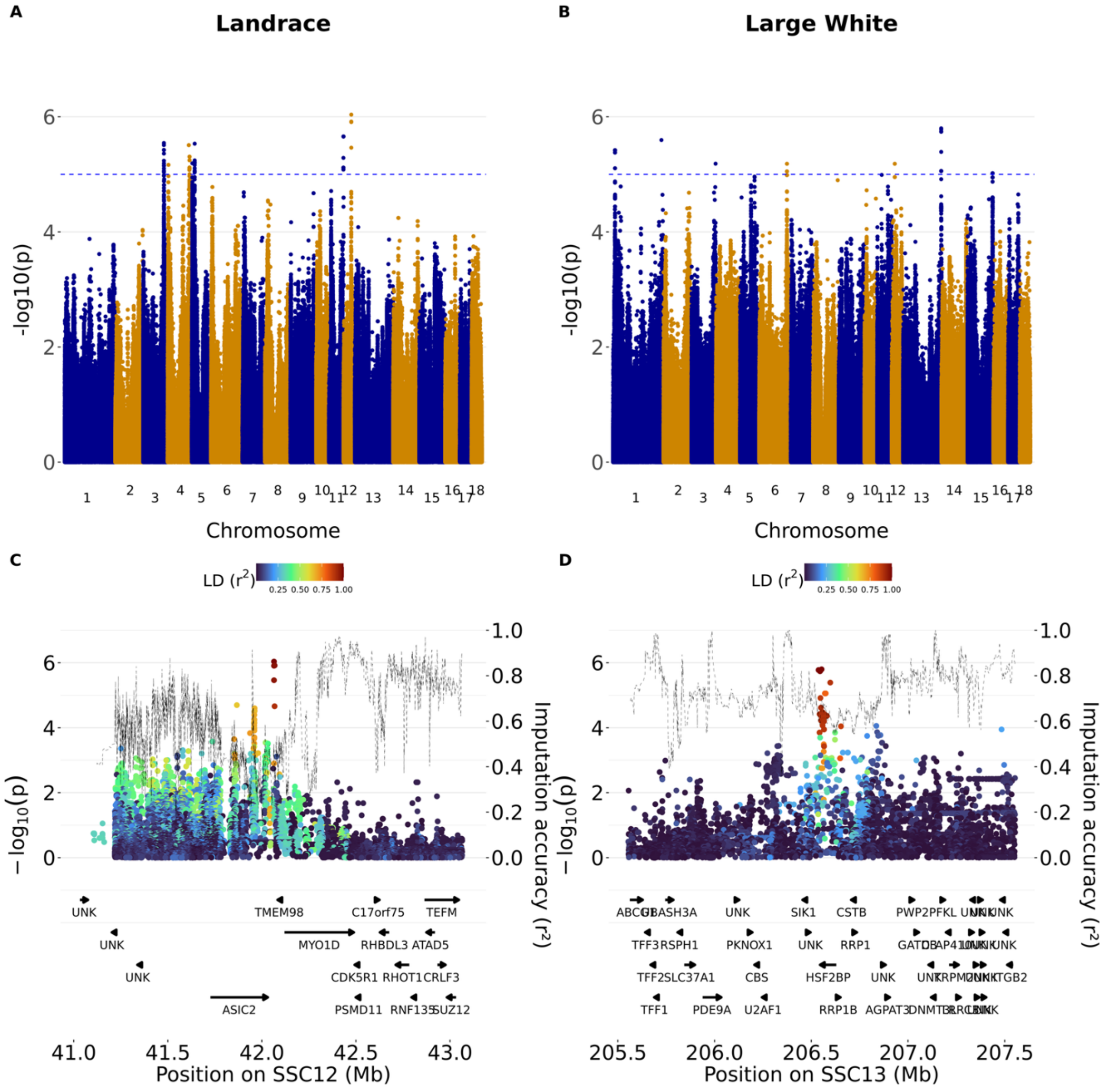


**Fig. S4. Genome-wide association results for the trait proportion of feed intake during the night ($FI_{night}$).**

Panels A and B show genome-wide association results for Swiss Landrace and Swiss Large White pigs. Panels C and D show regional association plots surrounding the lead variants. Dashed lines indicate genome-wide and suggestive significance thresholds. Colors indicate linkage disequilibrium ($r^2$) with the lead variant, and the dashed black line represents imputation accuracy.

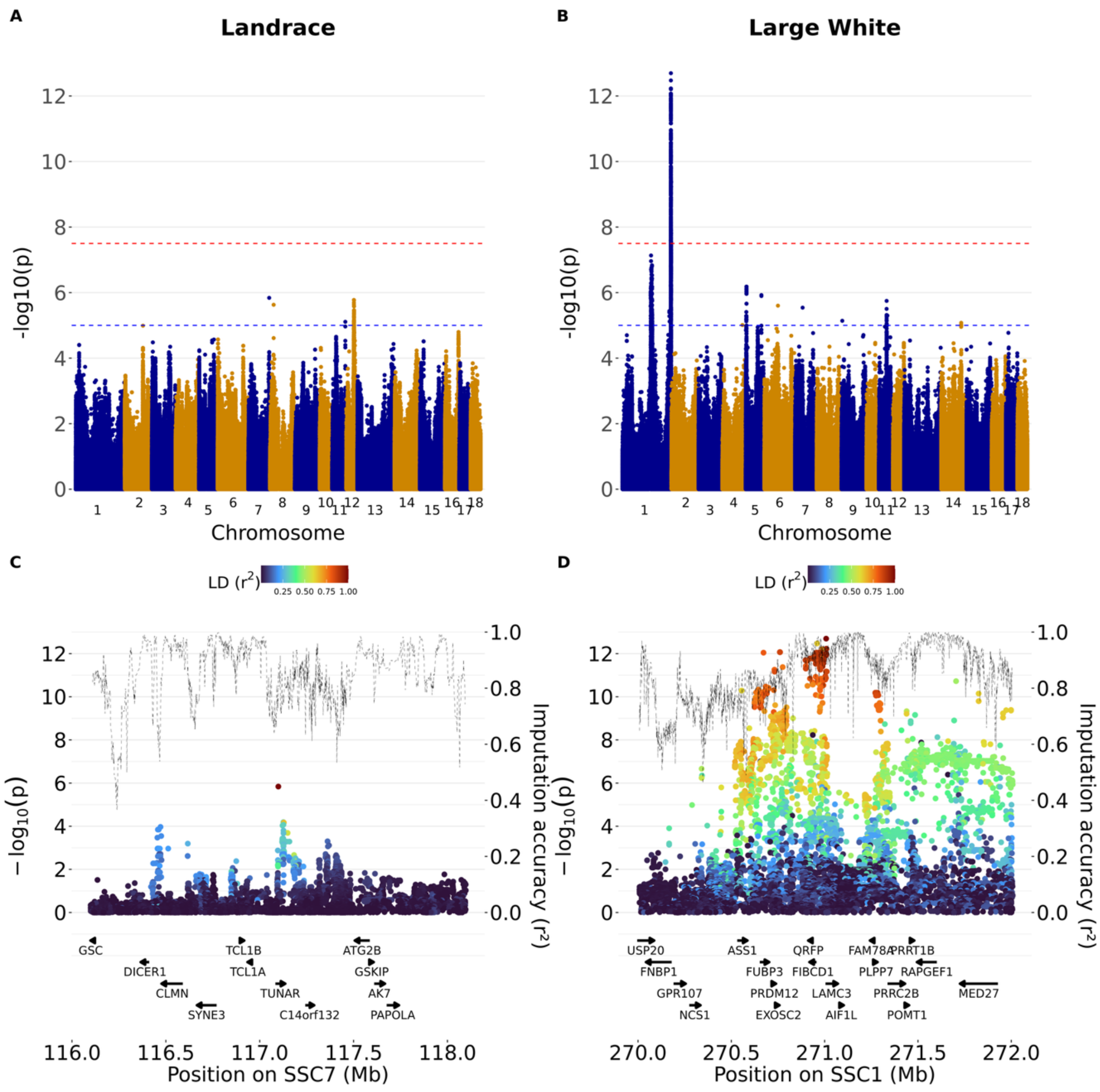


**Fig. S5. Genome-wide association results for the trait average daily gain (ADG).**

Panels A and B show genome-wide association results for Swiss Landrace and Swiss Large White pigs. Panels C and D show regional association plots surrounding the lead variants. Dashed lines indicate genome-wide and suggestive significance thresholds. Colors indicate linkage disequilibrium ($r^2$) with the lead variant, and the dashed black line represents imputation accuracy.

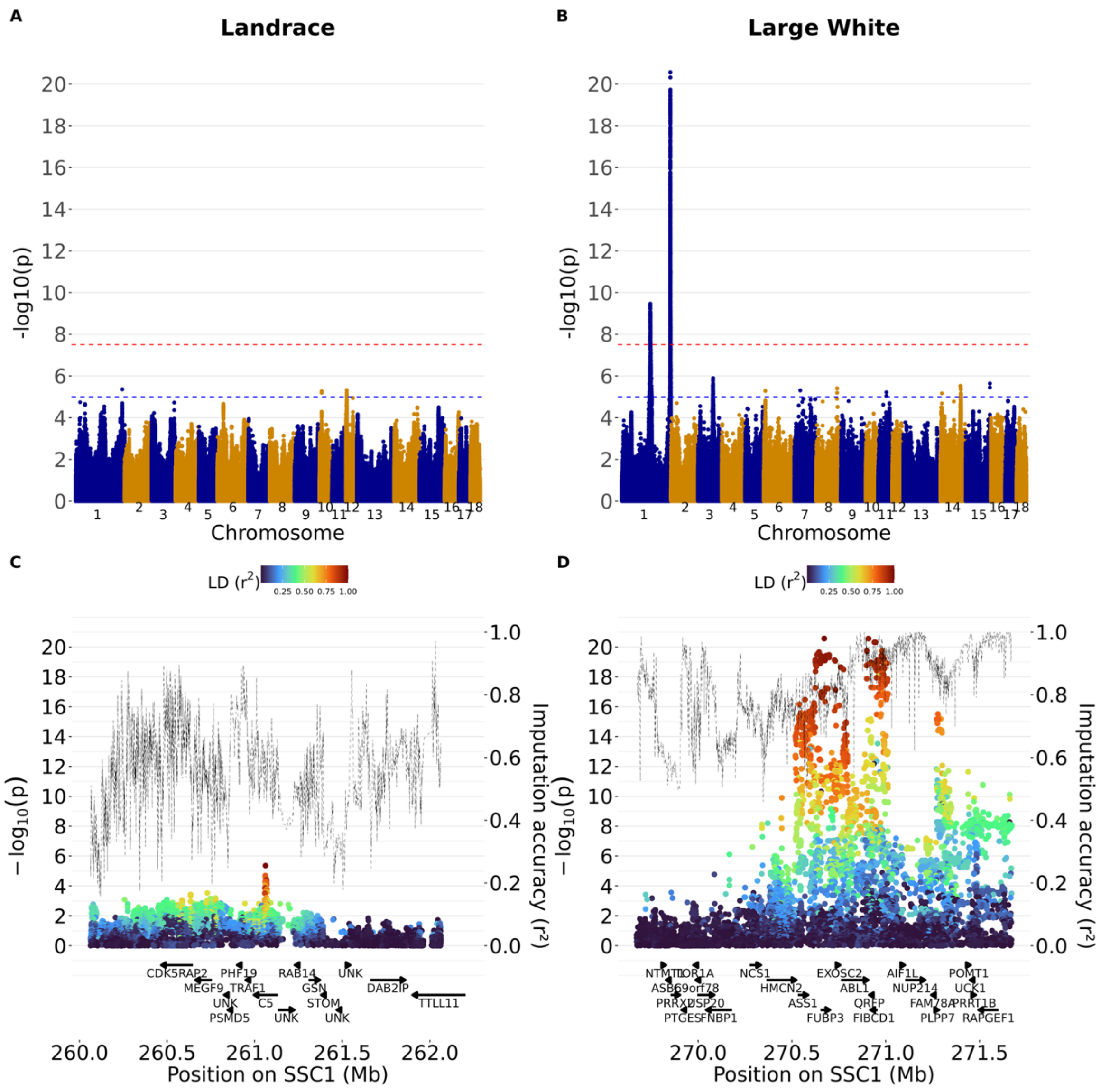


**Fig. S6. Genome-wide association results for the trait average daily feed intake (FI).**

Panels A and B show genome-wide association results for Swiss Landrace and Swiss Large White pigs. Panels C and D show regional association plots surrounding the lead variants. Dashed lines indicate genome-wide and suggestive significance thresholds. Colors indicate linkage disequilibrium (r²) with the lead variant, and the dashed black line represents imputation accuracy.

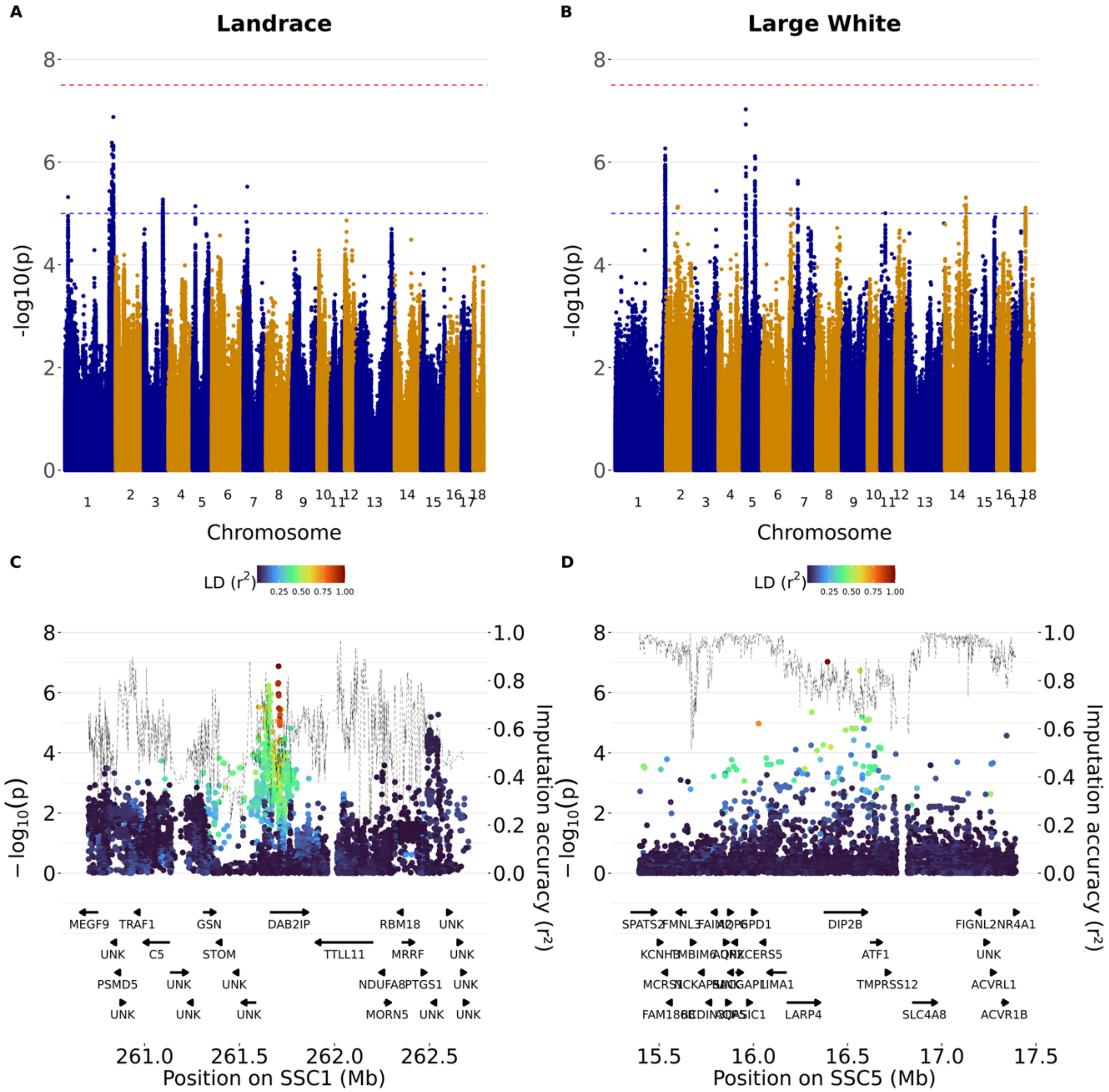


**Fig. S7. Genome-wide association results for the trait feed conversion ratio (FCR).**

Panels A and B show genome-wide association results for Swiss Landrace and Swiss Large White pigs. Panels C and D show regional association plots surrounding the lead variants. Dashed lines indicate genome-wide and suggestive significance thresholds. Colors indicate linkage disequilibrium (r²) with the lead variant, and the dashed black line represents imputation accuracy.

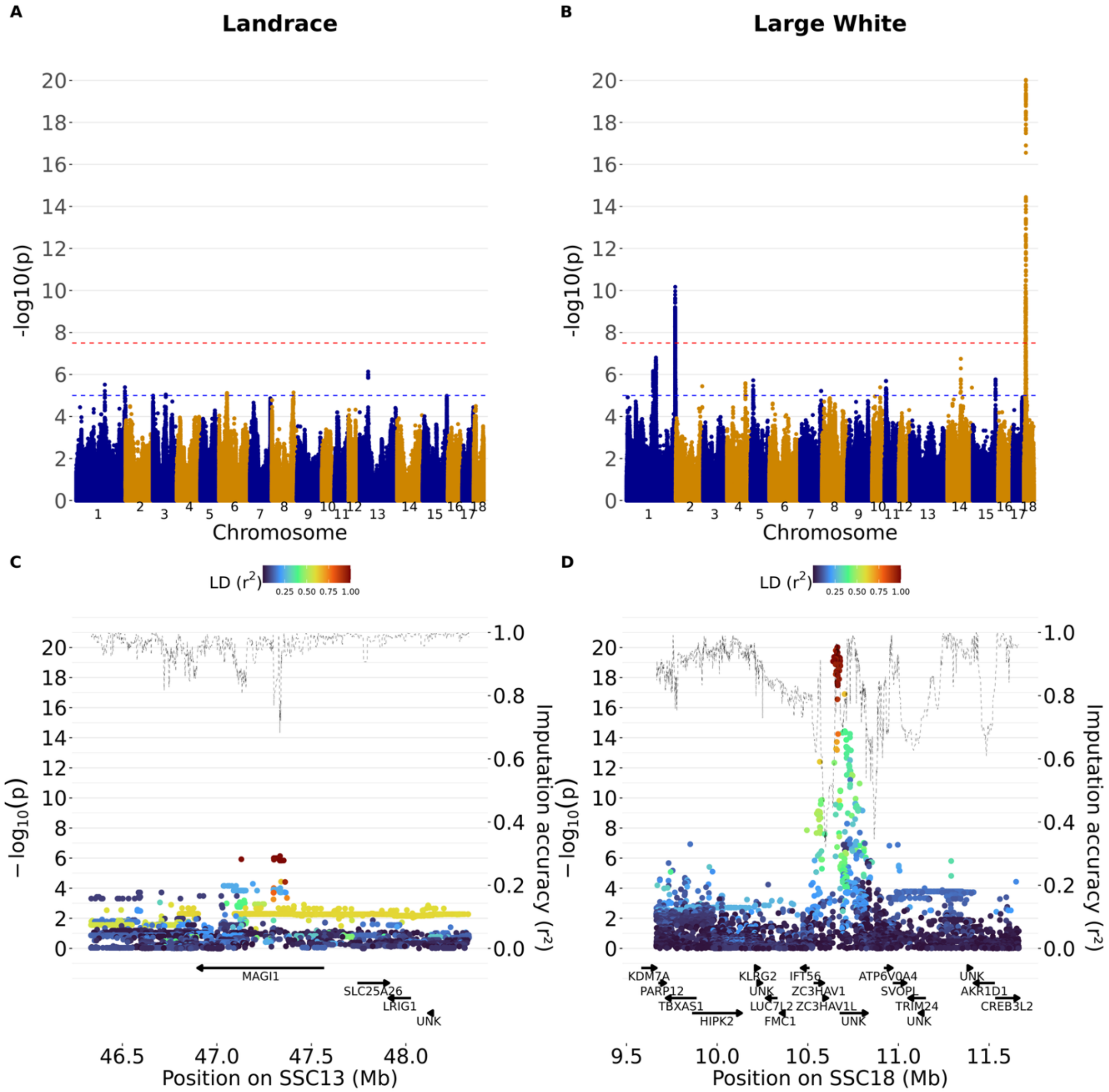


**Fig. S8. Genome-wide association results for the trait lean meat content (LMC).**

Panels A and B show genome-wide association results for Swiss Landrace and Swiss Large White pigs. Panels C and D show regional association plots surrounding the lead variants. Dashed lines indicate genome-wide and suggestive significance thresholds. Colors indicate linkage disequilibrium ($r^2$) with the lead variant, and the dashed black line represents imputation accuracy.

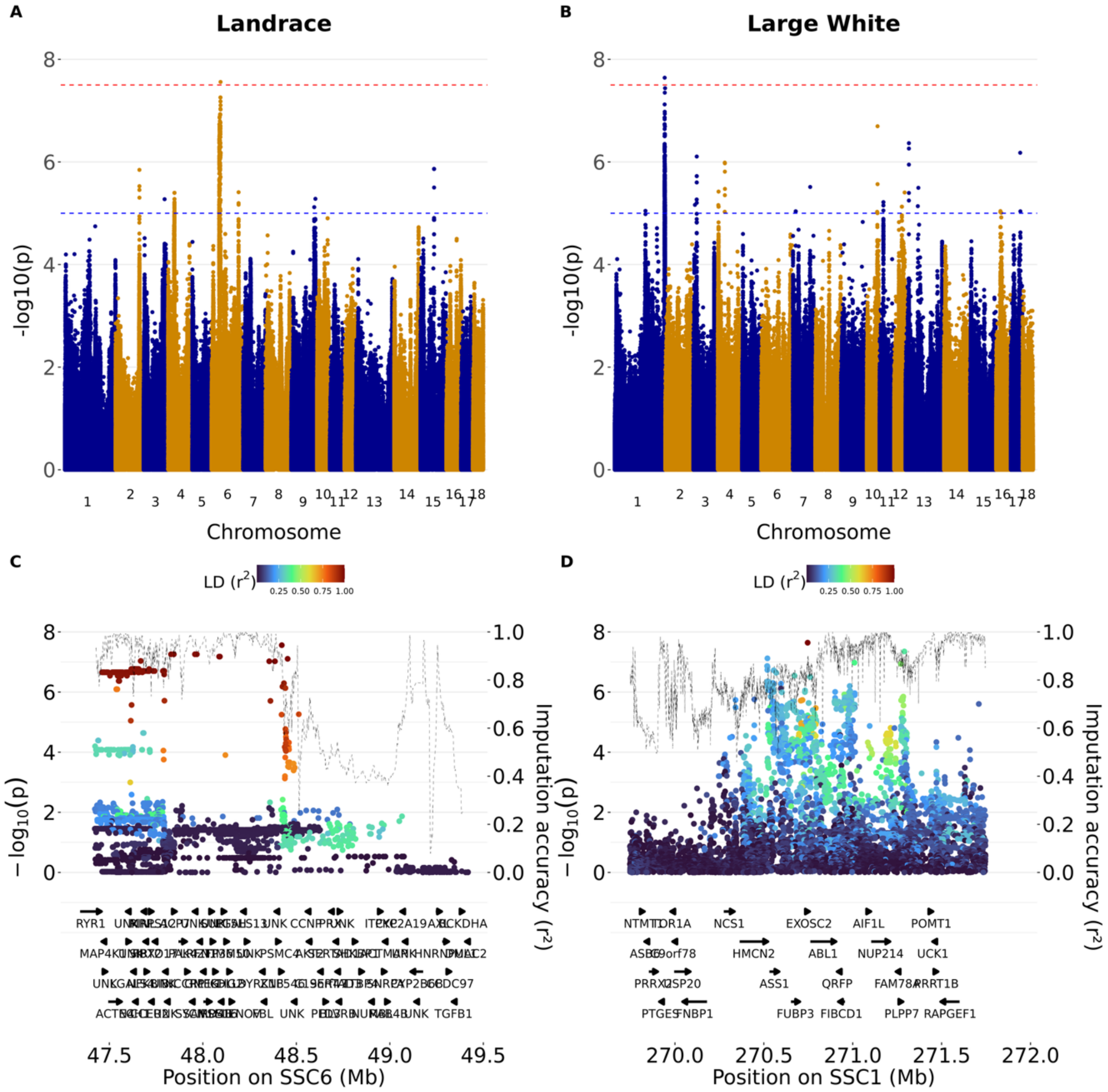


**Fig. S9. Genome-wide association results for the trait intramuscular fat (IMF).**

Panels A and B show genome-wide association results for Swiss Landrace and Swiss Large White pigs. Panels C and D show regional association plots surrounding the lead variants. Dashed lines indicate genome-wide and suggestive significance thresholds. Colors indicate linkage disequilibrium ($r^2$) with the lead variant, and the dashed black line represents imputation accuracy.

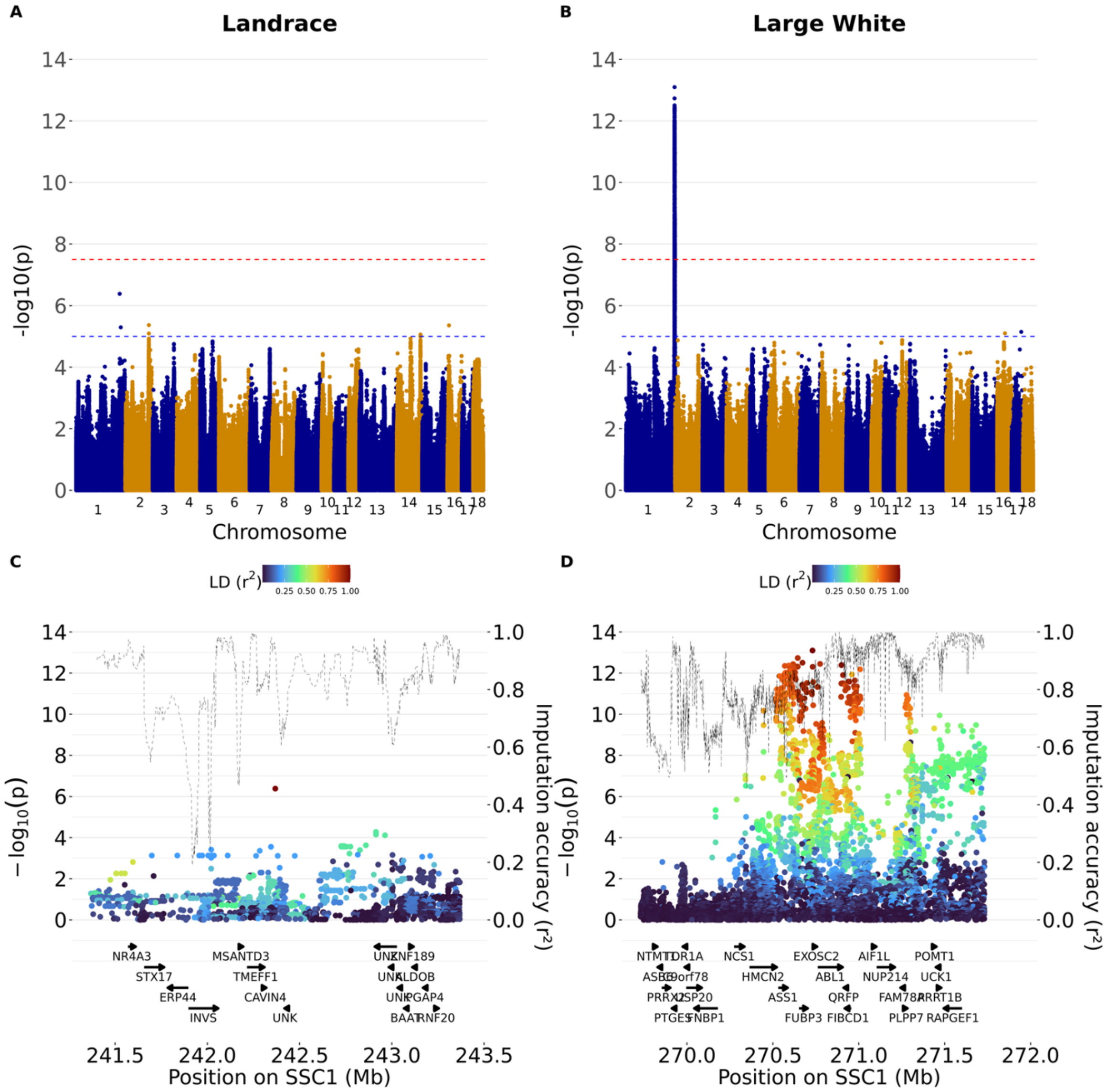


**Fig. S10. Genome-wide association results for the trait average daily feeding duration (DUR).**

Panels A and B show genome-wide association results for Swiss Landrace and Swiss Large White pigs. Panels C and D show regional association plots surrounding the lead variants. Dashed lines indicate genome-wide and suggestive significance thresholds. Colors indicate linkage disequilibrium ($r^2$) with the lead variant, and the dashed black line represents imputation accuracy.

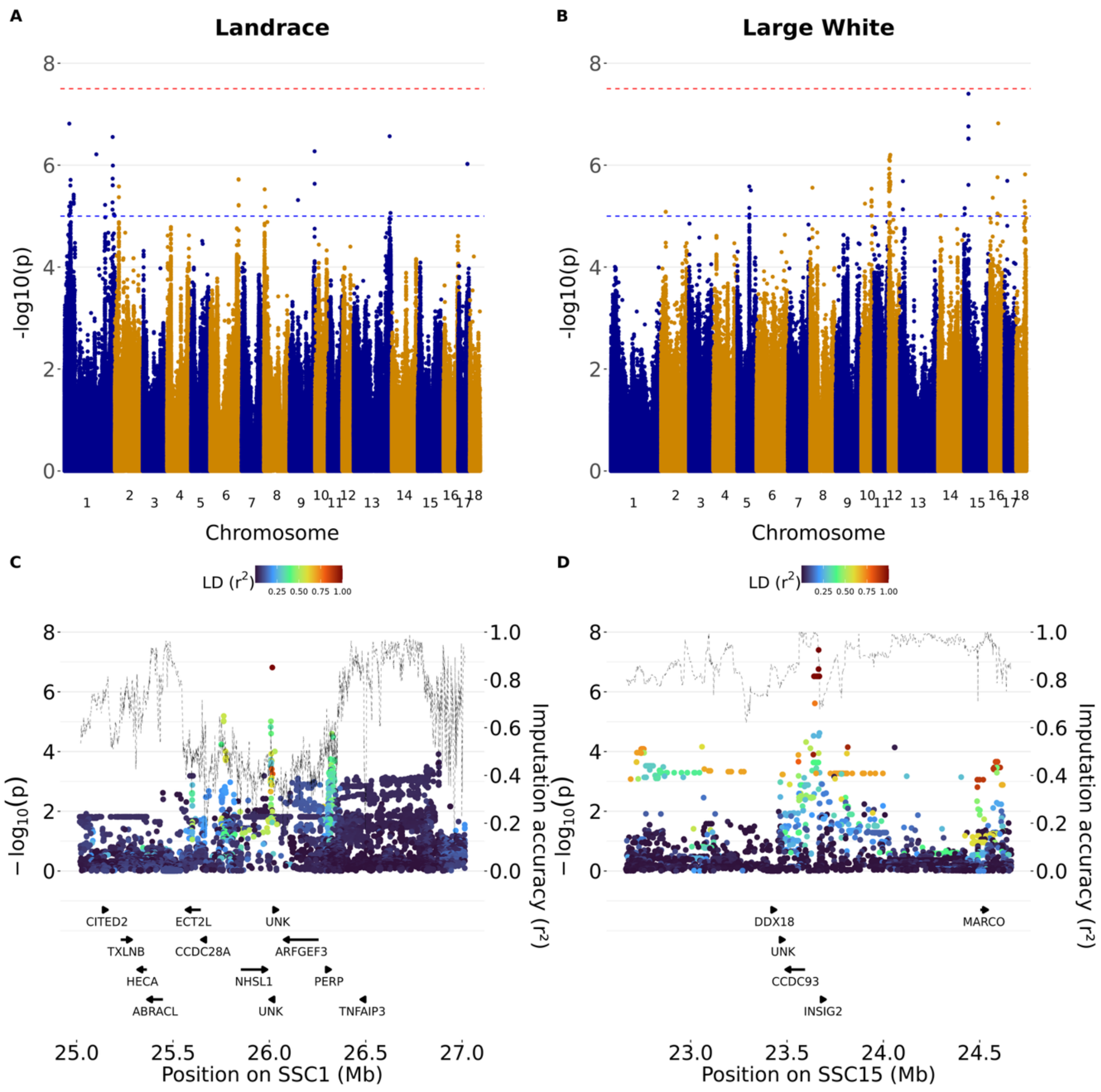


**Fig. S11. Genome-wide association results for the trait average daily feeding frequency (FREQ).**

Panels A and B show genome-wide association results for Swiss Landrace and Swiss Large White pigs. Panels C and D show regional association plots surrounding the lead variants. Dashed lines indicate genome-wide and suggestive significance thresholds. Colors indicate linkage disequilibrium ($r^2$) with the lead variant, and the dashed black line represents imputation accuracy.

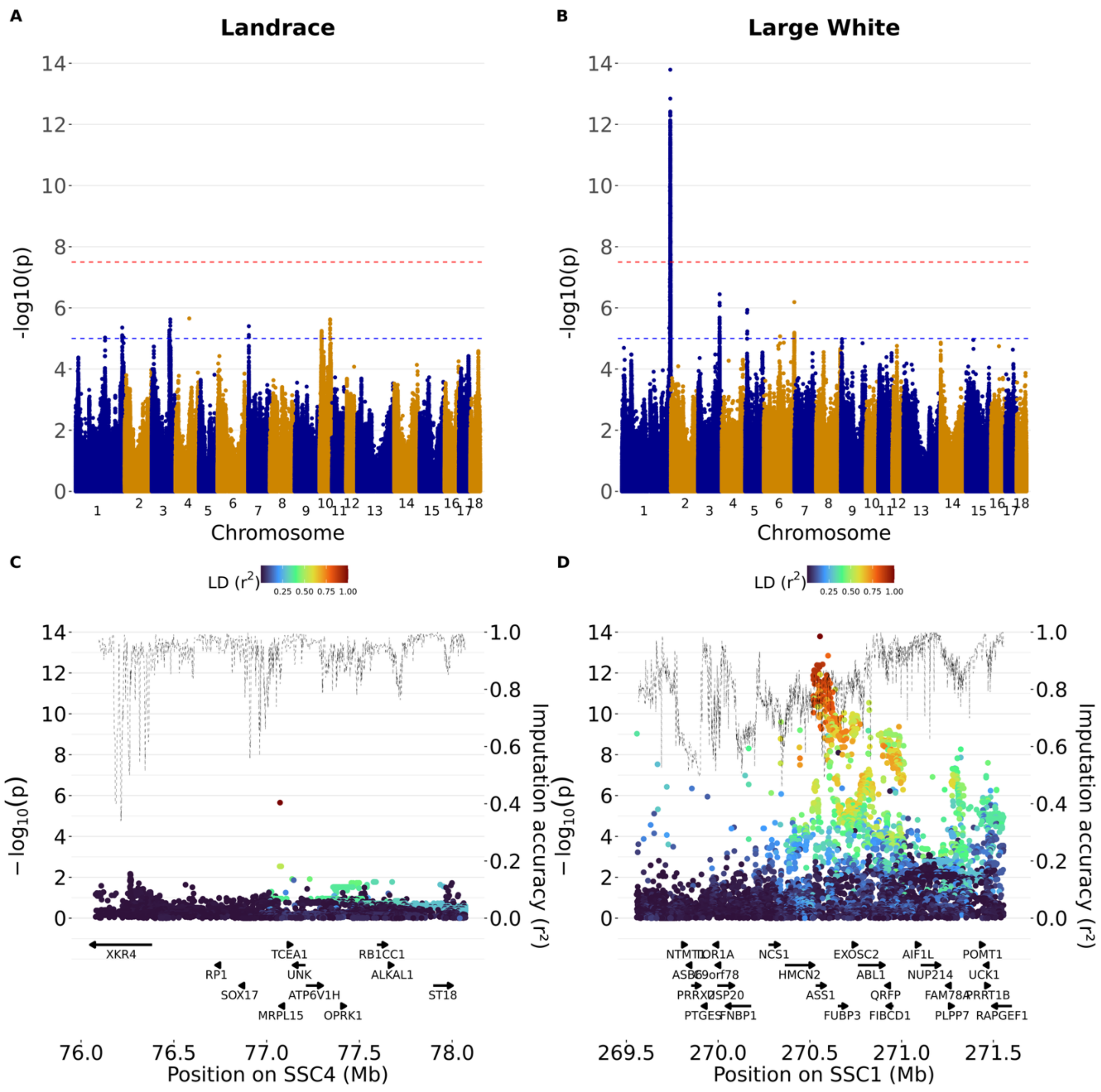


**Fig. S12. Genome-wide association results for the trait natural logarithm of variability in feed intake (Invar$_{FI}$).**

Panels A and B show genome-wide association results for Swiss Landrace and Swiss Large White pigs. Panels C and D show regional association plots surrounding the lead variants. Dashed lines indicate genome-wide and suggestive significance thresholds. Colors indicate linkage disequilibrium (r²) with the lead variant, and the dashed black line represents imputation accuracy.

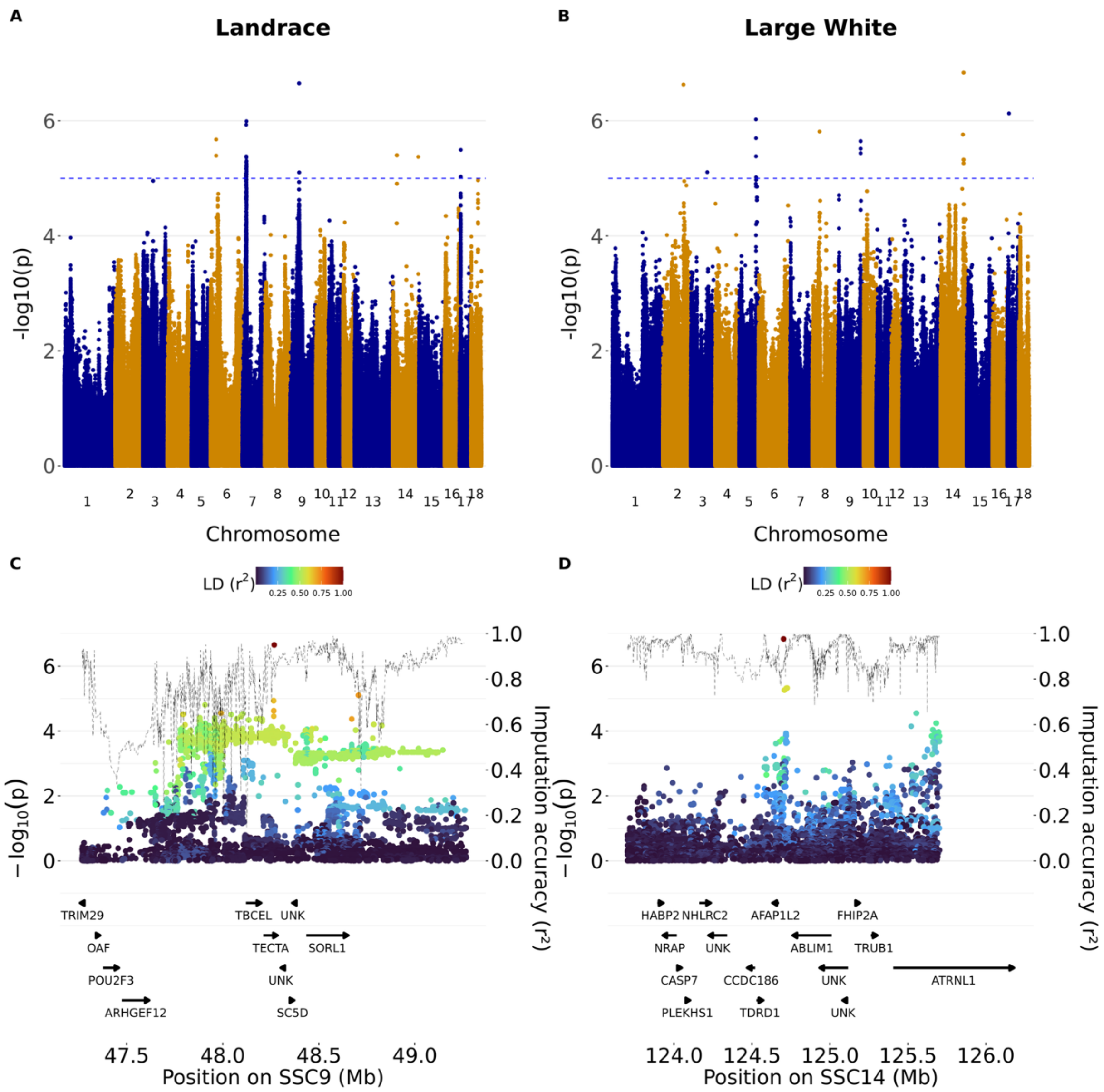


**Fig. S13. Genome-wide association results for the trait total number of medical treatments (TRT).**

Panels A and B show genome-wide association results for Swiss Landrace and Swiss Large White pigs. Panels C and D show regional association plots surrounding the lead variants. Dashed lines indicate genome-wide and suggestive significance thresholds. Colors indicate linkage disequilibrium ($r^2$) with the lead variant, and the dashed black line represents imputation accuracy.

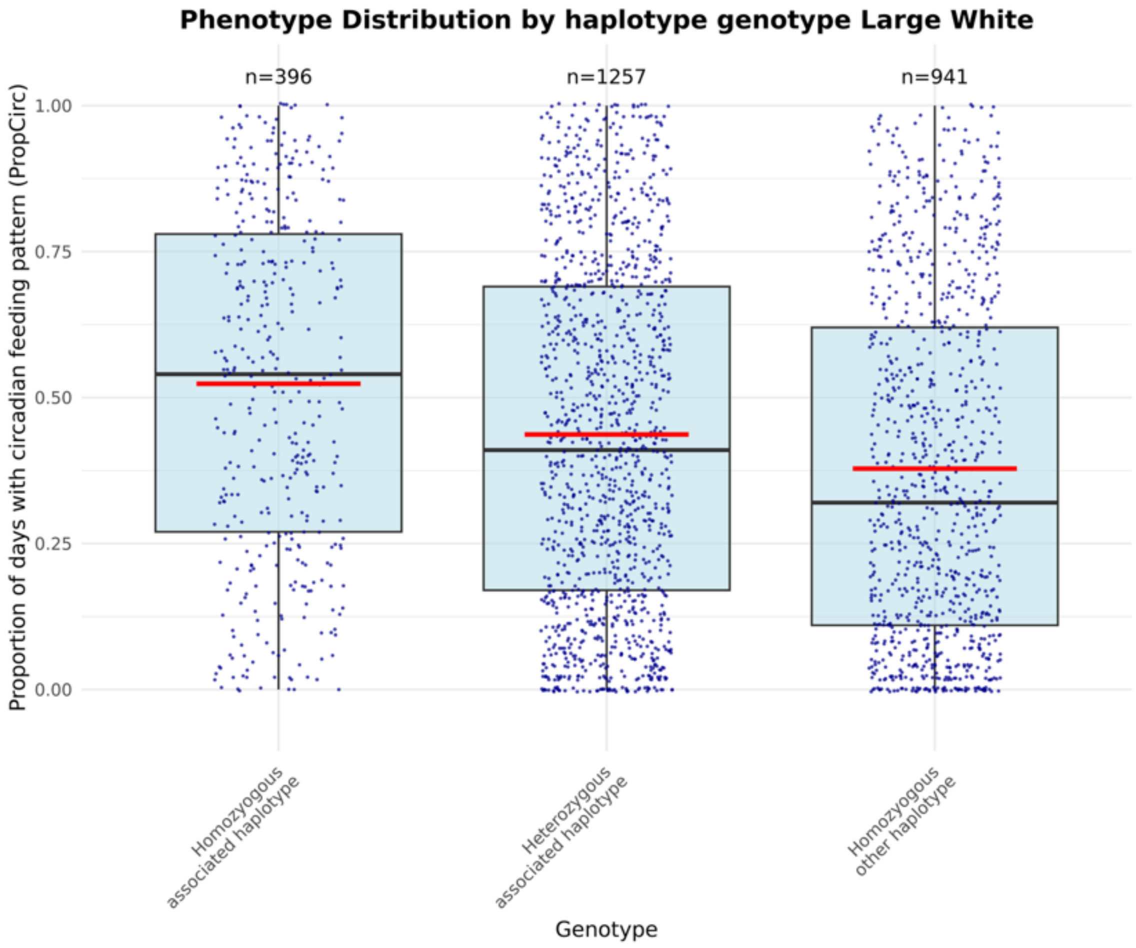


**Fig. S14. Effect of the lead haplotype on circadian feeding rhythmicity in Swiss Large White pigs.**

Boxplots showing PropCirc by lead haplotype genotype in Swiss Large White pigs.

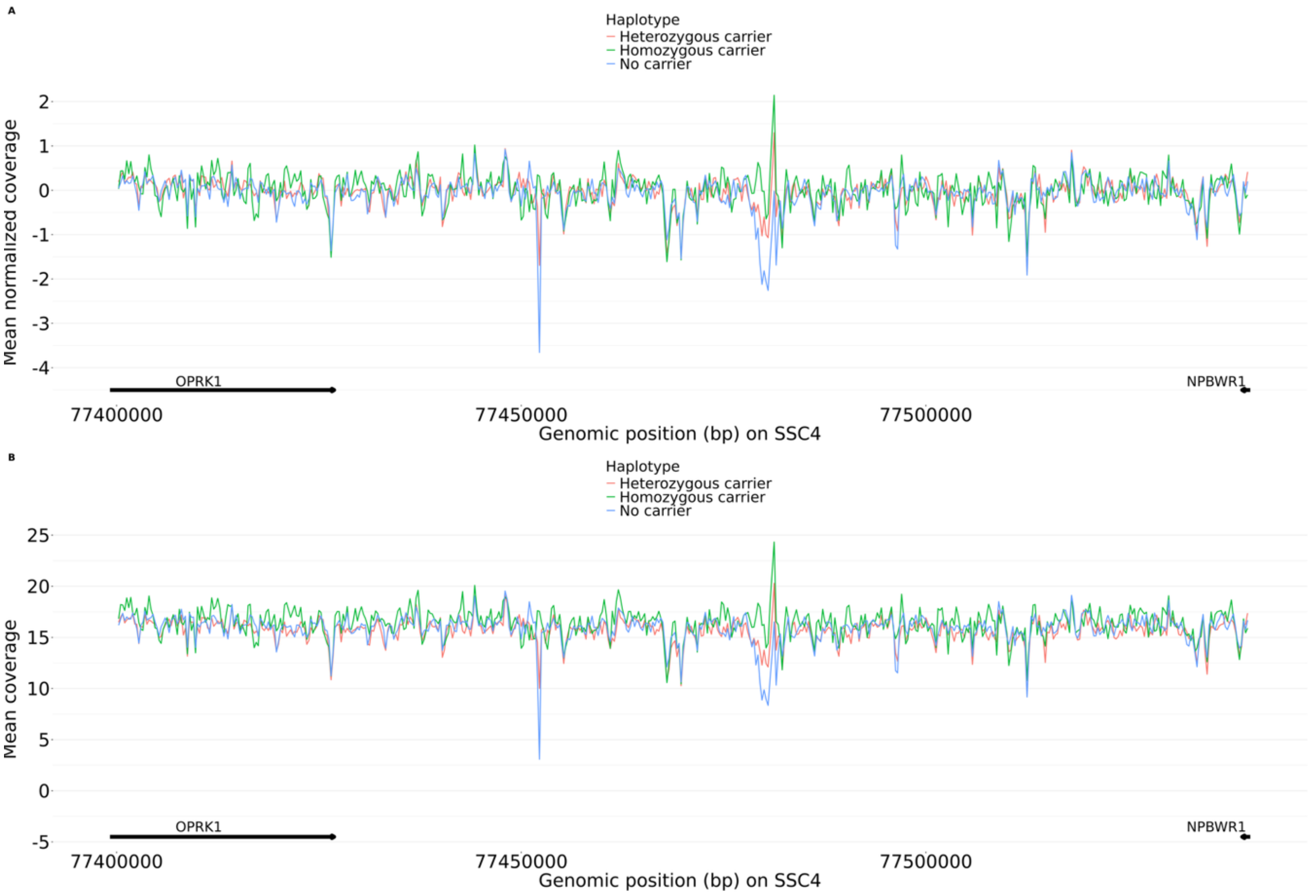


**Fig. S15. Sequence coverage differences associated with the top haplotype in Swiss Large White pigs.** Mean normalized coverage (A) and mean read coverage per 250 bp window (B) across the region surrounding the lead haplotype. A 1000bp region had a lower normalized coverage in homozygous haplotype carriers (Chr4:78,225,250-78,226,250; -3.7 normalized coverage), whereas a 250bp region located in between *OPRK1* and *NPBWR1* had a higher normalized coverage in homozygous haplotype carriers (Chr4:77,452,250-77,452,500; +4.0 normalized coverage), suggesting a deletion in non-carriers. A more exhaustive analysis of structural variants in eighteen pigs for which HiFi long-read sequencing data were available, confirmed a -307 bp deletion at Chr4: 77,452,166 in perfect LD with the top haplotype.

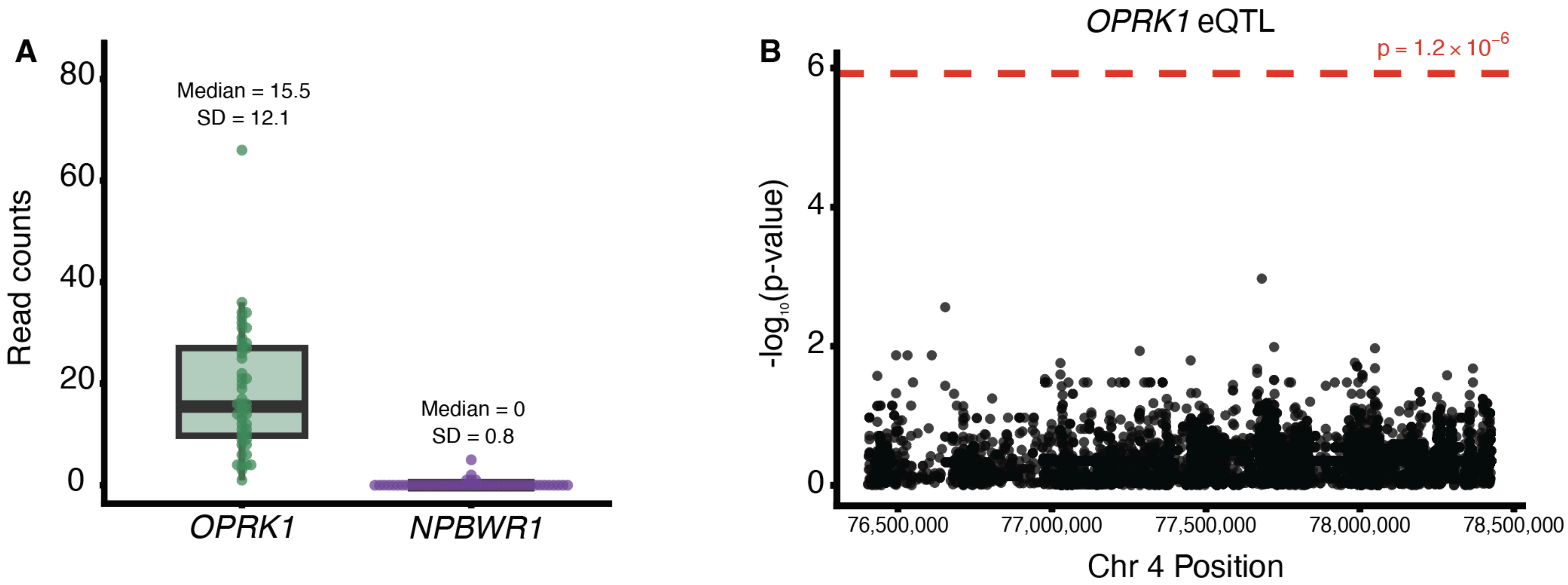


**Fig. S16. Expression of *OPRK1* and *NPBWR1* in 48 SLW hypothalamic samples and an eQTL analysis for *OPRK1.***

**(A)** Boxplot of the gene expression (read counts) for *OPRK1* and *NPBWR1*. Median read count and the standard deviation is listed above each gene. **(B)** Manhattan plot of the eQTL analysis for *OPRK1*, where the dotted red line represents the significance threshold. No genetic variants were associated with *OPRK1* expression.

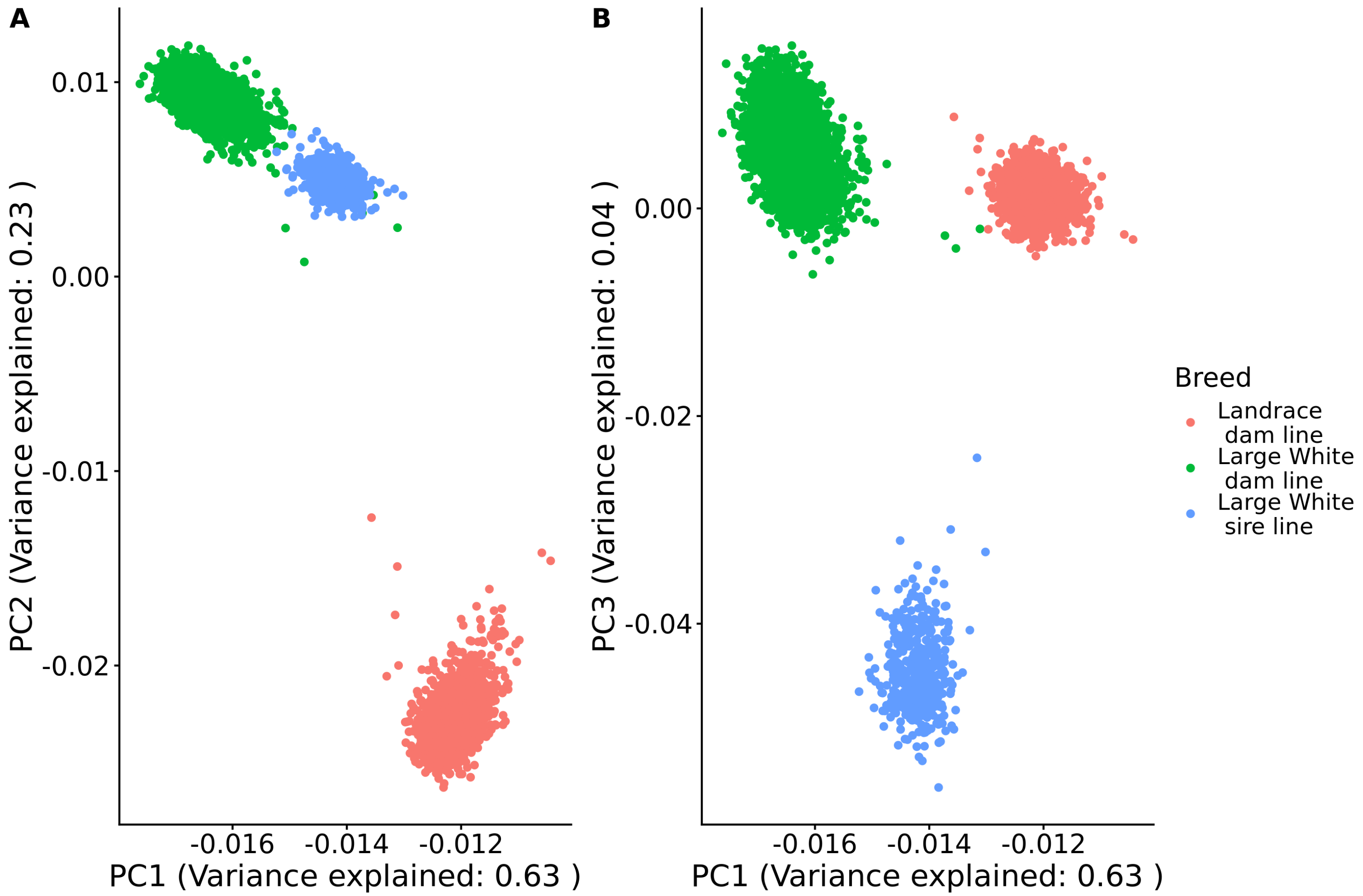


**Fig. S17. Principal component analysis of array-derived genotypes in Swiss Landrace and Swiss Large White pigs.**

Principal component analysis (PCA) based on SNP array genotypes. **(A)** PC1 vs. PC2; Swiss Large White samples form a distinct cluster. **(B)** PC1 vs. PC3.

#### Tables

**Table S1. Variance components and heritability estimates for circadian feeding and production traits in Swiss Landrace pigs.**

Legend:

Variance components were estimated using Bayesian animal models fitted with a single-step genomic relationship matrix. For each trait, posterior mean estimates and 95% highest posterior density (HPD) intervals are reported for additive genetic variance, residual variance, common environmental (group/pen) variance, and heritability ($h^2$).

Abbreviations: $FI_{night}$: Proportion of feed intake during night, ADG: Average daily gain, FI: Average daily feed intake, FCR: Feed conversion ratio, LMC: Lean meat content, DUR: Mean daily visit duration, FREQ: Mean daily visit frequency, $Invar_{FI}$: Feed intake resilience trait, TRT: number of medical treatments, IMF: intramuscular fat

| Trait | Genetic parameter | Mean estimate | HPD 95_min | HPD 95_max |
|---|---|---|---|---|
| PropCirc | Genetic Variance | 0.052 | 0.0426 | 0.0627 |
| PropCirc | Residual Variance | 0.0312 | 0.0263 | 0.0364 |
| PropCirc | Group Variance | 0.0084 | 0.0055 | 0.0121 |
| PropCirc | Heritability | 0.5668 | 0.4931 | 0.6343 |
| $FI_{night}$ | Genetic Variance | 0.0013 | 0.001 | 0.0015 |
| $FI_{night}$ | Residual Variance | 0.0008 | 0.0007 | 0.001 |
| $FI_{night}$ | Group Variance | 0.0003 | 0.0002 | 0.0004 |
| $FI_{night}$ | Heritability | 0.5217 | 0.4485 | 0.5886 |
| ADG | Genetic Variance | 779.6741 | 620.7 | 961.2 |
| ADG | Residual Variance | 556.6858 | 477.3 | 648.6 |
| ADG | Group Variance | 260.991 | 193.5 | 333.9 |
| ADG | Heritability | 0.4871 | 0.4022 | 0.5671 |
| FI | Genetic Variance | 0.0196 | 0.0147 | 0.0247 |
| FI | Residual Variance | 0.0227 | 0.02 | 0.0255 |
| FI | Group Variance | 0.0054 | 0.0036 | 0.0074 |
| FI | Heritability | 0.4098 | 0.3264 | 0.4853 |
| FCR | Genetic Variance | 0.0168 | 0.013 | 0.0209 |
| FCR | Residual Variance | 0.0152 | 0.0129 | 0.0173 |
| FCR | Group Variance | 0.0038 | 0.0026 | 0.0054 |
| FCR | Heritability | 0.4688 | 0.3826 | 0.5444 |
| LMC | Genetic Variance | 3.4715 | 3.021 | 3.994 |
| LMC | Residual Variance | 1.1721 | 0.9498 | 1.373 |
| LMC | Group Variance | 0.388 | 0.2414 | 0.5407 |
| LMC | Heritability | 0.6892 | 0.6326 | 0.7463 |
| IMF | Genetic Variance | 0.2291 | 0.1935 | 0.2691 |
| IMF | Residual Variance | 0.1467 | 0.1291 | 0.1662 |
| IMF | Group Variance | 0.0133 | 0.0047 | 0.0221 |
| IMF | Heritability | 0.5879 | 0.5275 | 0.6499 |
| DUR | Genetic Variance | 209669.327 | 175300 | 247300 |
| DUR | Residual Variance | 104561.097 | 88400 | 122300 |
| DUR | Group Variance | 44398.5536 | 32590 | 58710 |
| DUR | Heritability | 0.5838 | 0.5167 | 0.6487 |
| FREQ | Genetic Variance | 12.7494 | 9.12 | 17.27 |
| FREQ | Residual Variance | 23.2826 | 20.84 | 25.84 |
| FREQ | Group Variance | 46.1988 | 40.8 | 53.03 |
| FREQ | Heritability | 0.155 | 0.113 | 0.2009 |
| $Invar_{FI}$ | Genetic Variance | 0.0491 | 0.0294 | 0.0719 |
| $Invar_{FI}$ | Residual Variance | 0.1164 | 0.1027 | 0.1303 |
| $Invar_{FI}$ | Group Variance | 0.0434 | 0.0333 | 0.0544 |

| | | | | |
|---|---|---|---|---|
| $Invar_{FI}$ | Heritability | 0.2341 | 0.1438 | 0.3202 |
| TRT | Genetic Variance | 0.4092 | 0.1617 | 0.7059 |
| TRT | Residual Variance | 4.4001 | 4.059 | 4.745 |
| TRT | Group Variance | 0.0698 | 0.0032 | 0.1995 |
| TRT | Heritability | 0.0836 | 0.034 | 0.1429 |

**Table S2. Variance components and heritability estimates for circadian feeding and production traits in Swiss Large White pigs.**

Legend:

Variance components were estimated using Bayesian animal models fitted with a single-step genomic relationship matrix. For each trait, posterior mean estimates and 95% highest posterior density (HPD) intervals are reported for additive genetic variance, residual variance, common environmental (group/pen) variance, and heritability ($h^2$).

Abbreviations: $FI_{night}$: Proportion of feed intake during night, ADG: Average daily gain, FI: Average daily feed intake, FCR: Feed conversion ratio, LMC: Lean meat content, DUR: Mean daily visit duration, FREQ: Mean daily visit frequency, $lnvar_{FI}$: Feed intake resilience trait, TRT: number of medical treatments, IMF: intramuscular fat

| Trait | Genetic parameter | Mean estimate | HPD 95_min | HPD 95_max |
|---|---|---|---|---|
| PropCirc | Genetic Variance | 0.0443 | 0.0393 | 0.0481 |
| PropCirc | Residual Variance | 0.0327 | 0.0305 | 0.0355 |
| PropCirc | Group Variance | 0.0064 | 0.0054 | 0.0076 |
| PropCirc | Heritability | 0.5307 | 0.4849 | 0.5657 |
| $FI_{night}$ | Genetic Variance | 0.0009 | 0.0008 | 0.001 |
| $FI_{night}$ | Residual Variance | 0.0008 | 0.0008 | 0.0009 |
| $FI_{night}$ | Group Variance | 0.0002 | 0.0002 | 0.0002 |
| $FI_{night}$ | Heritability | 0.4689 | 0.4159 | 0.5127 |
| ADG | Genetic Variance | 585.6691 | 515.2 | 652.2 |
| ADG | Residual Variance | 542.9052 | 501.4 | 584.2 |
| ADG | Group Variance | 200.788 | 173.8 | 225.9 |
| ADG | Heritability | 0.4404 | 0.3967 | 0.48 |
| FI | Genetic Variance | 0.0198 | 0.018 | 0.0217 |
| FI | Residual Variance | 0.0124 | 0.0113 | 0.0136 |
| FI | Group Variance | 0.0055 | 0.0047 | 0.0062 |
| FI | Heritability | 0.5247 | 0.4873 | 0.5628 |
| FCR | Genetic Variance | 0.0164 | 0.0148 | 0.0181 |
| FCR | Residual Variance | 0.0113 | 0.0103 | 0.0123 |
| FCR | Group Variance | 0.0036 | 0.0031 | 0.0041 |
| FCR | Heritability | 0.5234 | 0.4846 | 0.5664 |
| LMC | Genetic Variance | 5.2444 | 2.319 | 11.39 |
| LMC | Residual Variance | 0.6919 | 0.0000 | 1.186 |
| LMC | Group Variance | 3.8751 | 0.0985 | 31.4 |
| LMC | Heritability | 0.6811 | 0.2665 | 0.9872 |
| IMF | Genetic Variance | 0.426 | 0.3887 | 0.461 |
| IMF | Residual Variance | 0.2356 | 0.2149 | 0.258 |
| IMF | Group Variance | 0.0148 | 0.0085 | 0.0221 |
| IMF | Heritability | 0.6296 | 0.5921 | 0.6658 |
| DUR | Genetic Variance | 261765.586 | 236600 | 285400 |
| DUR | Residual Variance | 156131.172 | 142100 | 169400 |
| DUR | Group Variance | 28148.1297 | 22650 | 34650 |
| DUR | Heritability | 0.5866 | 0.5437 | 0.6256 |
| FREQ | Genetic Variance | 41.2532 | 37.84 | 44.58 |
| FREQ | Residual Variance | 23.3128 | 21.58 | 25.24 |
| FREQ | Group Variance | 31.6198 | 29.1 | 34.44 |
| FREQ | Heritability | 0.4289 | 0.3985 | 0.4588 |
| $Invar_{FI}$ | Genetic Variance | 0.0351 | 0.0275 | 0.0431 |
| $Invar_{FI}$ | Residual Variance | 0.1184 | 0.1125 | 0.1239 |
| $Invar_{FI}$ | Group Variance | 0.0319 | 0.028 | 0.0362 |

| | | | | |
|---|---|---|---|---|
| $Invar_{FI}$ | Heritability | 0.1892 | 0.1511 | 0.229 |
| TRT | Genetic Variance | 0.3591 | 0.2061 | 0.5272 |
| TRT | Residual Variance | 4.7638 | 4.576 | 4.947 |
| TRT | Group Variance | 0.2551 | 0.1659 | 0.3445 |
| TRT | Heritability | 0.0667 | 0.039 | 0.0974 |

**Table S3. Genetic covariances and correlations between circadian feeding rhythmicity and production traits in Swiss Landrace pigs.**

Legend

Genetic parameters were estimated using bivariate Bayesian animal models. Reported values include posterior mean estimates and 95% highest posterior density (HPD) intervals for trait-specific variances, genetic covariance, and genetic correlation. Additional columns report Markov chain diagnostics including autocorrelation, effective sample size (ESS), total iterations, burn-in, and thinning interval.

Abbreviations: $FI_{night}$: Proportion of feed intake during night, ADG: Average daily gain, FI: Average daily feed intake, FCR: Feed conversion ratio, LMC: Lean meat content, DUR: Mean daily visit duration, FREQ: Mean daily visit frequency, $Invar_{FI}$: Feed intake resilience trait, TRT: number of medical treatments, IMF: intramuscular fat

See Excel file Table S3

**Table S4. Genetic covariances and correlations between circadian feeding rhythmicity and production traits in Swiss Large White pigs.**

Legend

Genetic parameters were estimated using bivariate Bayesian animal models. Reported values include posterior mean estimates and 95% highest posterior density (HPD) intervals for trait-specific variances, genetic covariance, and genetic correlation. Additional columns report Markov chain diagnostics including autocorrelation, effective sample size (ESS), total iterations, burn-in, and thinning interval.

Abbreviations: $FI_{night}$: Proportion of feed intake during night, ADG: Average daily gain, FI: Average daily feed intake, FCR: Feed conversion ratio, LMC: Lean meat content, DUR: Mean daily visit duration, FREQ: Mean daily visit frequency, $Invar_{FI}$: Feed intake resilience trait, TRT: number of medical treatments, IMF: intramuscular fat

See Excel file Table S4

**Table S5. Genome partitioning of phenotypic and genetic variance for circadian feeding rhythmicity in Swiss Landrace pigs.**

Legend

Genetic Variance components were estimated jointly using 19 genomic relationship matrices (GRMs), with one GRM for each autosome except chromosome 4, which was partitioned into a 5-Mb QTL region centered on the breed-specific lead variant and the remainder of chromosome 4. The chromosome 4 QTL region spanned 75,097,154–80,097,154 bp and was centered on the lead variant at Chr4:77,597,154. For each component, the variance estimate and standard error are reported. Phenotypic variance explained was calculated as $V(G_i)/V_P$, whereas genetic variance explained was calculated as $V(G_i)/\sum V(G)$. Residual variance V(e), total SNP-based genetic variance, and total phenotypic variance $V_P$ are also reported.

| GCTA label | Region | Variance component | SE | Phenotypic variance explained V(component)/Vp | SE | Genetic variance explained |
|---|---|---|---|---|---|---|
| V(G1) | Chr1 | 0.003972 | 0.001684 | 4.8% | 2.0% | 9.4% |
| V(G2) | Chr2 | 0.002912 | 0.001334 | 3.5% | 1.6% | 6.9% |
| V(G3) | Chr3 | 0.003177 | 0.001539 | 3.9% | 1.8% | 7.5% |
| V(G18) | Chr4 remainder | 0.000664 | 0.000999 | 0.8% | 1.2% | 1.6% |
| **V(G19)** | **Chr4 QTL region** | **0.004332** | **0.002644** | **5.3%** | **3.1%** | **10.3%** |
| V(G4) | Chr5 | 0.002230 | 0.001306 | 2.7% | 1.6% | 5.3% |
| V(G5) | Chr6 | 0.002886 | 0.001361 | 3.5% | 1.6% | 6.8% |
| V(G6) | Chr7 | 0.001679 | 0.001094 | 2.0% | 1.3% | 4.0% |
| V(G7) | Chr8 | 0.002072 | 0.001214 | 2.5% | 1.5% | 4.9% |
| V(G8) | Chr9 | 0.001919 | 0.001240 | 2.3% | 1.5% | 4.5% |
| V(G9) | Chr10 | 0.001598 | 0.001069 | 1.9% | 1.3% | 3.8% |
| V(G10) | Chr11 | 0.000409 | 0.000777 | 0.5% | 0.9% | 1.0% |
| V(G11) | Chr12 | 0.000386 | 0.000693 | 0.5% | 0.8% | 0.9% |
| V(G12) | Chr13 | 0.004995 | 0.001857 | 6.1% | 2.2% | 11.8% |

| | | | | | | |
|---|---|---|---|---|---|---|
| V(G13) | Chr14 | 0.003257 | 0.001440 | 4.0% | 1.7% | 7.7% |
| V(G14) | Chr15 | 0.001017 | 0.000965 | 1.2% | 1.2% | 2.4% |
| V(G15) | Chr16 | 0.002325 | 0.001117 | 2.8% | 1.3% | 5.5% |
| V(G16) | Chr17 | 0.001066 | 0.000910 | 1.3% | 1.1% | 2.5% |
| V(G17) | Chr18 | 0.001322 | 0.000953 | 1.6% | 1.2% | 3.1% |
| **V(e)** | **Residual variance** | **0.040187** | **0.002648** | **48.8%** | **4.2%** | |
| **Total genetic** | **Sum of V(G)** | | | **51.2%** | **4.2%** | **100.0%** |
| **Vp** | **Total phenotypic variance** | **0.082406** | **0.004505** | **100.0%** | | |

**Table S6. Genome partitioning of phenotypic and genetic variance for circadian feeding rhythmicity in Swiss Large White pigs.**

Legend

Genetic Variance components were estimated jointly using 19 genomic relationship matrices (GRMs), with one GRM for each autosome except chromosome 4, which was partitioned into a 5-Mb QTL region centered on the breed-specific lead variant and the remainder of chromosome 4. The chromosome 4 QTL region spanned 74,987,163–79,987,163 bp and was centered on the SLW lead variant at Chr4:77,487,163. For each component, the variance estimate and standard error are reported. Phenotypic variance explained was calculated as $V(G_i)/V_P$, whereas genetic variance explained was calculated as $V(G_i)/\sum V(G)$. Residual variance V(e), total SNP-based genetic variance, and total phenotypic variance $V_P$ are also reported.

| GCTA label | Region | Variance component | SE | Phenotypic variance explained V(component)/Vp | SE | Genetic variance explained |
|---|---|---|---|---|---|---|
| V(G1) | Chr1 | 0.004422 | 0.001310 | 5.6% | 1.6% | 12.2% |
| V(G2) | Chr2 | 0.004672 | 0.001395 | 5.9% | 1.7% | 12.9% |
| V(G3) | Chr3 | 0.001850 | 0.000812 | 2.3% | 1.0% | 5.1% |
| V(G18) | Chr4 remainder | 0.001575 | 0.000860 | 2.0% | 1.1% | 4.3% |
| **V(G19)** | **Chr4 QTL region** | **0.002313** | **0.001121** | **2.9%** | **1.4%** | **6.4%** |
| V(G4) | Chr5 | 0.001589 | 0.000798 | 2.0% | 1.0% | 4.4% |
| V(G5) | Chr6 | 0.003438 | 0.001093 | 4.3% | 1.4% | 9.5% |
| V(G6) | Chr7 | 0.000653 | 0.000592 | 0.8% | 0.7% | 1.8% |
| V(G7) | Chr8 | 0.000811 | 0.000610 | 1.0% | 0.8% | 2.2% |
| V(G8) | Chr9 | 0.000864 | 0.000677 | 1.1% | 0.9% | 2.4% |
| V(G9) | Chr10 | 0.000937 | 0.000685 | 1.2% | 0.9% | 2.6% |
| V(G10) | Chr11 | 0.001550 | 0.000747 | 2.0% | 0.9% | 4.3% |
| V(G11) | Chr12 | 0.001731 | 0.000782 | 2.2% | 1.0% | 4.8% |
| V(G12) | Chr13 | 0.000426 | 0.000532 | 0.5% | 0.7% | 1.2% |
| V(G13) | Chr14 | 0.003082 | 0.001036 | 3.9% | 1.3% | 8.5% |

| | | | | | | |
|---|---|---|---|---|---|---|
| V(G14) | Chr15 | 0.003083 | 0.001041 | 3.9% | 1.3% | 8.5% |
| V(G15) | Chr16 | 0.001859 | 0.000854 | 2.3% | 1.1% | 5.1% |
| V(G16) | Chr17 | 0.000610 | 0.000582 | 0.8% | 0.7% | 1.7% |
| V(G17) | Chr18 | 0.000771 | 0.000599 | 1.0% | 0.8% | 2.1% |
| **V(e)** | **Residual variance** | **0.043107** | **0.001978** | **54.3%** | **3.2%** | |
| **Total genetic** | **Sum of V(G)** | | | **45.7%** | **3.2%** | **100.0%** |
| **Vp** | **Total phenotypic variance** | **0.079342** | **0.002919** | **100.0%** | | |

**Table S7. eQTL within 1 Mb of the top PropCirc GWAS variant for SLW**

Legend

Gene: the gene name; Top eQTL position: location of each gene's most significant eQTL along Chromosome 4; Top eQTL nominal p-value: nominal p-value of the most significant variant for each gene; Number of variants tested: number of variants considered for each gene's eQTL analysis; Significance threshold: significance threshold determined by the beta-distribution after 1,000 permutations and an FDR of 5%. NA indicates a failure to establish a beta threshold for that gene; Distance from SLW GWAS variant: distance between the location of the most significant variant in the SLW GWAS and the gene's transcription start site; SLW GWAS variant p-value: nominal p-value from the eQTL analysis for the most significant variant in the SLW GWAS.

| Gene | Top eQTL position | Top eQTL nominal p-value | Significance threshold | Number of variants tested | Distance from SLW lead GWAS variant | SLW GWAS variant p-value |
|---|---|---|---|---|---|---|
| ENSSSCG00000046734 | 76,057,697 | 0.000150936 | 1.27869e-05 | 9,809 | 897,436 | 0.857652 |
| ENSSSCG00000056807 | 76,282,861 | 1.79623e-05 | NA | 11,562 | 721,234 | 0.328924 |
| *SOX17* | 76,333,777 | 0.00176593 | 1.39025e-06 | 12,810 | 630,015 | 0.993523 |
| *MRPL15* | 77,468,330 | 0.000615377 | 1.18901e-06 | 15,150 | 403,440 | 0.898642 |
| *RGS20* | 76,493,894 | 0.000884627 | 2.18861e-06 | 16,135 | 277,630 | 0.341464 |
| ENSSSCG00000055986 | 77,359,235<br>77,359,231 | 0.00286378 | 1.49175e-06 | 18,979 | 107,059 | 0.2454690 |
| ENSSSCG00000057277 | 78,633,819 | 0.00309205 | 1.12811e-06 | 21,124 | 607,080 | 0.167964 |
| ENSSSCG00000060401 | 77,155,273<br>77,311,510<br>77,367,944<br>77,452,516<br>77,640,622<br>79,018,046<br>79,018,067<br>79,059,661 | 0.00156745 | 5.58781e-07 | 21,137 | 611,664 | 0.473861 |

| | | | | | | |
|---|---|---|---|---|---|---|
| | 79,063,117<br>79,069,349 | | | | | |
| *PCMTD1* | 79,090,549<br>79,103,136<br>79,103,363 | 0.000449374 | 7.99918e-07 | 21,709 | 611,837 | 0.583739 |
| *ST18* | 78,869,227 | 0.00175402 | 3.06345e-07 | 20,835 | 410,575 | 0.228325 |
| ENSSSCG00000060320 | 77,487,163 | 4.36341e-05 | 5.50738e-07 | 20,123 | 381,742 | 0.233884 |
| ENSSSCG00000057329 | 78,468,394 | 0.00171905 | 6.80796e-08 | 19,938 | 261,544 | 0.331907 |
| ENSSSCG00000054250 | 77,704,523 | 0.00227893 | 4.60829e-07 | 19,748 | 268,829 | 0.0554614 |
| ENSSSCG00000051846 | 77,704,523 | 0.000665987 | 9.22227e-07 | 19,699 | 264,663 | 0.503823 |
| ENSSSCG00000051386 | 78,037,610<br>78,037,621 | 9.48941e-05 | 1.64059e-07 | 19,508 | 163,704 | 0.0341544 |
| *RB1CC1* | 77,717,234 | 0.000272313 | 2.02776e-06 | 19,497 | 107,162 | 0.529567 |
| *OPRK1* | 77,681,425 | 0.00106778 | 1.19992e-06 | 17,919 | 87,979 | 0.238969 |
| *ATP6V1H* | 76,487,709 | 0.00203653 | 1.71825e-06 | 17,333 | 275,669 | 0.443899 |
| *TCEA1* | 76,501,926<br>76,516,890<br>76,527,828<br>76,743,731<br>77,026,712 | 0.00208936 | 4.50121e-07 | 15,565 | 377,568 | 0.341030 |
| *LYPLA1* | 77,083,549 | 0.000573624 | 3.38570e-07 | 15,247 | 403,614 | 0.137762 |
| ENSSSCG00000045551 | 75,934,309<br>76,023,592<br>77,444,448<br>77,456,882<br>77,467,987<br>77,468,038 | 0.000145489 | NA | 12,922 | 626,803 | 0.360968 |
| ENSSSCG00000060890 | 76,225,178 | 0.000214235 | 6.24657e-07 | 12,793 | 641,694 | 0.421408 |

**Dataset S1 (separate file). Genome-wide significant variants associated with circadian feeding rhythmicity in Swiss Landrace and Swiss Large White pigs.**

Legend

This dataset contains all SNPs reaching genome-wide significance ($P < 5 \times 10^{-8}$) in genome-wide association analyses for circadian feeding rhythmicity (PropCirc) in Swiss Landrace and Swiss Large White pigs. For each variant, the dataset reports chromosome, genomic position (bp), top SNP identifier, p-value, minor allele frequency (MAF), effect and reference alleles (A1/A2), and breed-specific association results. Analyses were performed using imputed whole-genome sequence variants. Only variants exceeding the genome-wide significance threshold in at least one breed are included.

See Excel File Dataset S1

**Dataset S2 (separate file). Functional annotation of genome-wide significant variants associated with circadian feeding rhythmicity using Ensembl VEP.**

Legend

This dataset contains functional annotations of genome-wide significant variants associated with circadian feeding rhythmicity (PropCirc) in Swiss Landrace and Swiss Large White pigs. Variants were annotated using the Ensembl Variant Effect Predictor (VEP). For each variant, the dataset reports genomic location, allelic information, predicted molecular consequence, functional impact, affected gene and transcript, and protein-level changes where applicable. Additional annotations include variant identifiers, existing database entries, and predictive scores where available. Only variants reaching genome-wide significance ($P < 5 \times 10^{-8}$) in the association analyses are included.

See Excel File Dataset S2
